# GuavaVision AI: An Explainable Deep Learning Framework for Automated Classification, Lesion Localization, and Segmentation of Guava Diseases

**DOI:** 10.64898/2026.06.18.733093

**Authors:** Jahanur Biswas, Muhaimenul Islam, Md. Maruf Bangabashi, Monira Akter, Tabia Sultana Nishi, MD Kawser Sheikh, Md. Rasel Mia, Md Musfique Anwar

**Affiliations:** Department of Computer Science and Engineering, Dhaka International University, Dhaka, Bangladesh; Department of Computer Science and Engineering, Jahangirnagar University, Dhaka, Bangladesh

## Abstract

Guava cultivation is considerably influenced by foliar and fruit diseases whose overlapping symptoms and environmental variability make accurate field-level diagnosis challenging. Numerous studies have been conducted to find efficient methods of diagnosing plant diseases, but most focus on image-level classification and do not include lesion localization or pixel-level segmentation of the images within a single framework of analysis. This study proposes a comprehensive framework for utilizing automated image analysis to classify guava leaf and fruit diseases at the image level, locate lesions, and segment lesions at the pixel level from multiple images of the same type of disease collected from various growing conditions. The dataset was enriched through three augmentation strategies including standard preprocessing, structured augmentation, and GAN-based synthetic image generation, expanding the effective training data to approximately 7,000 images, while a 5-fold cross-validation strategy guided model selection and final performance was assessed on a held-out test set. The experimental evaluation of multiple state-of-the-art Convolutional Neural Networks (CNNs) for the classification of guava leaf and fruit diseases indicated that the model generated using the ResNet50+DenseNet121 model fusion achieved the highest classification accuracy of 98.20%. For lesion detection and segmentation, YOLOv8-seg outperformed Mask R-CNN, achieving mAP@0.5 of 0.907 and 0.889, and mAP@0.5:0.95 of 0.783 and 0.769 for detection and segmentation, respectively, with a balanced precision–recall profile. The techniques of Explainable AI (XAI) were used to increase the transparency of this model by identifying areas in the image that are significant to the actual lesion. The framework was further designed with practical web-based deployment in mind, evaluating both lightweight and high-capacity models to balance computational efficiency against predictive accuracy. From this research, it was concluded that using model fusion, data augmentation, and segmentation-aware lesion detection would provide a solution for managing guava diseases effectively.

## 1. Introduction

The guava (*Psidium guajava*) is a widely grown plant throughout the tropics and subtropical regions. Due to its nutritional value and ability to adapt to a wide variety of growing environments, it is also commonly referred to as “Queen of Fruits” [1]. Guavas are a good source of potassium, dietary fiber, antioxidants, vitamin C, and support both immune system function and cardiovascular health [2]. Guava is also an important part of the fruit production system in developing countries because of its ability to adapt to many different climates [3], and it has a strong economic importance in these areas because of its contributions to local economies.

Even with the economic importance of guava, production is susceptible to numerous illnesses induced by foliar and fruit fungal, bacterial, and viral pathogens [3]. These diseases usually lead to decreased yield quality, increased production costs, and increased dependence on chemical treatments. Disease outbreak costs have the potential to be very high in major guava-producing areas, such as Bangladesh, where guava is produced on a total of approximately 0.3 million hectares per year with a total production of over 3.7 million metric tons [4]. In addition, many factors, such as variations in cultivar type, growth stage, and environmental conditions, frequently result in similar visual symptoms, making reliable field-level diagnosis challenging.

Farmers and agricultural specialists typically use only visual assessment methods to diagnose diseases. Nonetheless, such evaluations are intrinsically subjective and may exhibit inconsistency, especially when symptom patterns overlap or vary with illumination and humidity conditions [5]. While there are reliable diagnostic tests, like those that utilize real-time polymerase chain reaction and mass spectrometry, they require expensive equipment and a person with technical expertise to operate [6]. Therefore, making the use of these systems impractical for large-scale disease identification efforts. These limitations have resulted in an increased interest in automated imaging systems for disease diagnosis using image-based plant analysis.

The advancement of machine learning and deep learning technology has produced automated analysis of plant disease with considerable results [6]. Research into the use of convolutional neural networks (CNN) for the classification of guava leaf diseases has reported both high rates of accuracy. As an example, the Inception-ResNet50 method achieved an accuracy of 97.50% [7], while there is also a reported accuracy of 97.32% using a multi-fusion meta-learning approach [8]. Additionally, the combination of EfficientNetV2 and Vision Transformers yielded an accuracy of approximately 95%, with the addition of interpretability via Grad-CAM [9]. Similarly, DenseNet201 has been reported to achieve an accuracy of 96% for classifying the fruit disease [10]. Other types of segmentation-based models (GLSM, YOLOv5, and GIP-MU-Net), which are aimed at localizing infected areas more precisely, have also been reported to have been explored beyond that of image classification [5].

Despite the above-mentioned studies involving the classification of guava leaf diseases using CNN and segmentation-based models, the vast majority of studies have employed only image-level prediction methods and do not combine lesion localization and pixel-level segmentation methods into a single framework [5–10]. As such, these existing methods are limited in their applicability for performing quantitative severity estimation and implementing targeted intervention. Furthermore, the vast majority of studies employed a limited number of images, as well as imbalanced datasets. There is little consistency in the use of cross-validation, nor has the interpretability of models been given the same importance as the physical attributes of models when designing and conducting research. Additionally, there are very few studies that evaluate detection and segmentation models in the same annotation and training conditions as it pertains to either of these methods; thus, fairness and accuracy of comparison are extremely difficult to achieve.

These considerations strongly imply that a single, segmentation-aware and interpretable framework is needed to balance both the rigor of the methodology as well as the practicality of the application for guava disease management.

The contributions of the current study are listed below:

1. Development of a standardized deep learning framework that includes the three parts of classification, localization of the lesion, and pixel-level segmentation under a unified evaluation system.
2. Implementation of a systematic approach to enhance the data for achieving better generalized models across different agricultural field environments.
3. Providing complete benchmarks for the performance of several Convolutional Neural Networks (CNNs), including a classification model developed using a fusion of images.
4. The extensive evaluation of the two state-of-the-art detection and segmentation networks (YOLOv8-Seg and Mask R-CNN) for the localization of lesions within an image and the estimation of severity based on those localized lesions.
5. Integration of Grad-CAM–based explainability and deployment of a lightweight web-based decision support system for use in agricultural applications.

## 2. Literature Review

Recent advancements in machine learning and deep learning techniques have dramatically improved automated detection of plant diseases through image analysis. Different convolutional neural network (CNN) architectures and segmentation models have been employed for diagnosing and detecting diseases on fruit and leaf samples. Several studies on classifying and detecting guava diseases have been conducted. Other fields of research on plant disease detection have resulted in the development of new CNN architectures and learning methods. This section indicates a large number of studies relevant to the proposed method and highlights where limitations exist, which necessitate the need for the proposed method.

A hybrid architecture combining GIP-MU-Net for infected patch segmentation, GLSM for leaf region segmentation, and a YOLOv5-based module for multi-disease detection from a single leaf was suggested by Rashid et al. [5]. A total of 2,542 guava leaf images were used in the dataset during evaluation. There were five disease categories or groups to classify these images. The accuracy of the segmentation stage of the framework was 92.41%. The detection stage of the framework achieved the following: 73.3% precision; 73.1% recall, and 71% mAP@0.5. Comparatively, the segmentation stage achieved better performance results than the detection stage due to using stricter IoU thresholds during segmentation than detection. Other research has focused on feasibility from the end-user perspective instead of multi-component frameworks, such as the device-friendly framework proposed by Nandi et al. [6]. In that work, quantization (float16, dynamic range quantization, etc.) was applied to decrease model size while maintaining comparable levels of accuracy for guava fruit and leaf diseases using a dataset of 1,834 images, achieving 94.93% classification accuracy. Guler et al. [7] explored ensemble and fusion methods for classifying guava diseases through an evaluation of deep networks under hybrid augmentation using 2,063 guava leaf images, where the InceptionV3–ResNet50 ensemble achieved 97.50% accuracy with an F1-score of 0.975 and AUC of 0.9934. In a related vein, Asim et al. [8] applied an ensemble machine-learning approach with multi-fusion metacognition on 560 guava leaf images across seven disease categories, obtaining 97.32% classification accuracy, which supports the idea that using feature-level fusion together with meta-learning can be useful for limited datasets.

In addition to building reliable models that accurately classify objects, an increasing number of recent research efforts consider additional factors such as explainability and deployment of results. Farooqui et al. [9] used a hybrid EfficientNetV2–Vision Transformer model, applied the Grad-CAM algorithm for highlighting disease-mediated areas of tissue, and created an online portal for users to access detection results and explanations; their model achieved 95% classification accuracy across five guava leaf disease classes. Researchers have explored guava fruits for an array of disease identification, but studies can also serve to inform fine-grained classification tasks. Toptas et al. [10] set out to classify guavas commercially using 2,310 fruit images representing nine classes, studying several varieties and ripening states while evaluating several pretrained CNN models through both stratified k-fold cross-validation and an 80:20 random train/test split. Comparison of the metrics obtained from models based on the architectural backbone of DenseNet201 showed the best performance with approximately 96.5% accuracy. In addition to using deep learning to conduct analysis of guavas, research has also employed classical machine learning approaches to analyze guava diseases within constrained data domains. For example, Oya et al. [11] applied preprocessing steps such as segmentation, contrast enhancement (CLAHE), and augmentation prior to training their models using 473 guava fruit images, where SVM and ANN classifiers achieved close to 99% accuracy. Their findings suggest that pipeline design and preprocessing steps can significantly impact results achieved within any model class.

Many areas of research involving plant diseases provide an enormous number of publications dealing with validation methods, choices of models, and the move to a more comprehensive analysis of lesions rather than simply identifying their presence. Using hybrid deep learning frameworks in detecting diseases on coffee leaves, Thakur et al. [12] proposed SUNet, which combines U-Net segmentation, SegNet encoding, VGG16 feature extraction, and Mask R-CNN instance segmentation. Experiments conducted on the JMuBEN and JMuBEN2 datasets demonstrated improved accuracy, IoU, precision, recall, and F1-score compared to baseline segmentation approaches. In order to detect apple leaf diseases, Parashar et al. [13] combined the Inception-v3 CNN architecture with 5-fold cross-validation on 1,821 apple leaf images, achieving 94.76% classification accuracy with balanced precision and recall. Transfer learning has also been investigated by Ç etiner et al. [14], who compared several pretrained architectures, including VGG19, DenseNet169, MobileNet, Xception, and NASNetLarge, on a dataset of 3,651 apple leaf images, achieving approximately 98% classification accuracy after systematic preprocessing. Integrated pipelines are emerging in the field of plant disease. Polly et al. [15] utilized a CNN-based classification system along with YOLOv8 detection and DeepLabV3+/U-Net segmentation to achieve an overall accuracy of 96.97%. Additionally, Umar et al. [16] combined CNN classification with YOLOv7 detection in order to classify the diseases of tomato leaves with an accuracy of 98.8%.

The size of datasets and deployment are often pragmatic restrictions to system design. Islam et al. [17] trained a series of deep learning models on a massive set of 10,000 leaf images. The ResNet50 model performed the best, with 98.6% accuracy, and the resulting model has been introduced into a web application. On the same note, Faisal et al. [18] augmented an existing citrus disease dataset of 759 images to 3,383 and indicated a testing accuracy of 99.48% with EfficientNetB3. Bezabh et al. [19] adopted a comparable approach to the detection of mango leaf disease, increasing their data set to 3,195 images and 1,452 to 3,195, and obtained 99.21% testing accuracy with a GoogLeNet-VGG ensemble, which indicates the benefits of using complementary feature extractors.

Studies that emphasize segmentation are promising in that they can provide important evidence that can be used to improve lesion delineation. Haider et al. [20] studied the EFFS-Net and MDFS-Net segmentation architectures using various agricultural datasets with the highest Dice coefficients (95.01), thus demonstrating that segmentation-based lesion detection could provide useful information to estimate the severity of the disease. Belay et al. [21] augmented a dataset of chickpea disease cases to 8,391 images using augmentation methods and suggested a CNN-LSTM model with the result of a classification rate of 92.55%. Madhurya et al. [22] proposed the YR2S deep-learning architecture based on YOLOv7 detections and hybrid feature aggregation and achieved a detection accuracy of 99.69%. More comparative studies have shown that the best architecture can change based on the type of data. Nagamani et al. [23] compared Fuzzy-SVM, CNN, and R-CNN models on tomato disease detection, but Rahman et al. [24] compared a variety of CNN architectures on a large multi-crop dataset with 30,945 images of 35 disease classes and a classification with an accuracy of 95.62%.

Recent studies on guava show that there is growing dependence on transfer learning, interpretability, and detection models to mimic orchard conditions. Ahmed et al. [25] transferred ResNet 101 to a guava leaf image dataset of 3,784 leaf images, which achieved a classification accuracy of 98.48%. Grad-CAM was used to confirm that the network focused on disease-affected areas. According to Chiou et al. [26], the YOLOv4 algorithm was applied to detect the defects on guava fruits, where 189 samples of guava fruits were transformed into 1,701 augmented images, and the detection rate was 88.15%. Mirjat et al. [27] studied guava wilt detection on 1,420 leaf images segmented using convolutional neural network models and achieved a classification accuracy of 95.17%, and emphasized the need to detect diseases early.

Transfer learning and hybrid architectures remain the most popular methodological practices among different crop species. Laur et al. [28] employed an EfficientNet-based transfer learning framework to identify diseases in grapevines, attaining an accuracy of approximately 98.7%. Arshad et al. [29], on the other hand, presented the PLDPNet hybrid architecture involving the combination of CNN ensembles with transformer-based feature fusion that demonstrated an accuracy of 98.66% and the F1-score of 96.33% when it comes to the detection of potato leaf diseases. Alataawi et al. [30] evaluated models of deep learning models in a comparative study with a dataset of 15,915 plant leaf images with 19 disease types and an accuracy of 95.2% using the VGG16 architecture. Also, Ali et al. [31] established that the use of class-weighting strategies and data augmentation significantly improves the performance of an EfficientNet-based classifier in the event of data imbalance.

The importance of detection and segmentation models in the context of real agricultural systems has been highlighted by recent research. Mamat et al. [32] used YOLOv5 on the dataset of about 3,500 images and categorized the fruit types with an accuracy of 99.5%. Yuan et al. [33] re-configured DeepLab v3+ for grape leaf lesion segmentation and obtained a mean intersection-over-union (mIoU) of 0.848 and F1-score of 0.918 on 1,180 grape leaf lesion images. Khan et al. [34] presented a comparative study of several variants of YOLO in detecting maize disease and found that YOLOv8n achieved the highest accuracy of 99.04%. To explain its predictions, Shafik et al. [35] incorporated SHAP-based explainable AI approaches to study the results of a deep-learning model using 4,447 images of plant diseases, achieving an accuracy of 97.41%.

It has also been noted that large benchmark datasets have made significant contributions to the advancement of research in the identification of plant diseases. Andrew J. et al. [36] tested various deep learning designs on the PlantVillage dataset of 54,305 images of 38 plant diseases, with DenseNet-121 having the highest accuracy of 99.81 per cent. Fruit quality has also been assessed using detection-based systems. The authors Mukhiddinov et al. [37] presented a better YOLOv4 model trained on 12,000 images of fruits and vegetables with a training accuracy of 75.8% and testing accuracy of 73.5% to classify fresh and rotten food. A VGG-19 transfer learning model with HSV segmentation was proposed by Thanh-Hai Nguyen et al. [38] to classify tomato disease on 16,010 images of the PlantVillage dataset with 99.72% accuracy, with shorter training time. Mimma et al. [39] suggested a pipeline of YOLOv3 followed by YOLOv7 to detect fruits and ResNet50/VGG16 to classify them (99 and 98 percent accuracy, respectively, on a self-generated dataset with eight classes of fruits). Zayani et al. [40] used YOLOv8 to detect tomato diseases, achieved a 66.67% accuracy with an augmented Roboflow dataset, and explored the capability of class imbalance to affect model performance. Nouman Butt et al. [41] presented a deep learning and Moth-Flame Optimization model on the detection of citrus diseases with an accuracy of 99.6% with the use of DenseNet-201 and AlexNet. U-Net was used in Elinisa et al. [42] to segment banana diseases on 18,240 images, with a Dice coefficient of 96.45 and IoU of 93.23. Lastly, Rijon et al. [43] have offered a CNN-Gradient Boosting ensemble in the classification of the guava disease, in which EfficientNet-B0 feature extraction was combined with Gradient Boosting classifiers, resulting in an accuracy of 99.99%.

A summary of the representative research in the field of crop disease detection and guava disease analysis is outlined in Table 1. Although deep learning methodologies have been shown to be highly effective at demonstrating high accuracy in image-based classification tasks, relatively few studies have addressed lesion detection and then followed by pixel-wise segmentation to realize an effective localization; indeed, most studies have largely focused on classification only. The use of relatively small datasets and heterogeneous evaluation protocols is also an additional limitation that could hinder the application of model results to the agronomic situations in the real world. In addition, there exist a few studies that combine explainable AI paradigms with overall diagnostic pipelines. These weaknesses emphasize the need to create integrated and interpretable frameworks that can address classification, detection, and pixel segmentation to enable an efficient analysis of plant diseases within a farming context.

**Table 1.** Summary of representative studies on plant and guava disease detection.

| Author (Ref.) | Crop | Task | Model / Method | Dataset Size | Key Result | Limitation |
| --- | --- | --- | --- | --- | --- | --- |
| Rashid et al. [5] | Guava | Segmentation + Detection | GIP-MU-Net + GLSM + YOLOv5 | 2542 images | Segmentation accuracy: 92.41% | Detection performance lower |
| Nandi et al. [6] | Guava | Classification | GoogLeNet + model quantization | 1834 images | Accuracy: 94.93% | Only classification considered |
| Guler et al. [7] | Guava | Classification | InceptionV3-ResNet50 Ensemble | 2063 images | Accuracy: 97.50% | No lesion localization |
| Asim et al. [8] | Guava | Classification | Meta-learning ensemble | 560 images | Accuracy: 97.32% | Small dataset |
| Toptas et al. [10] | Guava | Fruit classification | DenseNet201 | 2310 images | Accuracy: 96.5% | Disease localization not addressed |
| Ahmed et al. [25] | Guava | Classification + XAI | ResNet101 + Grad-CAM | 3784 images | Accuracy: 98.48% | Image-level classification only |
| Chiou et al. [26] | Guava | Detection | YOLOv4 | 1701 images | Accuracy: 88.15% | Limited dataset size |
| Andrew J. et al. [36] | Multi-crop | Classification | DenseNet-121 | 54,305 images | Accuracy: 99.81% | Controlled dataset conditions |
| Mukhiddinov et al. [37] | Multi-crop | Detection | Improved YOLOv4 | 12,000 images | Precision: 75.8% | Lower testing precision |
| Thanh-Hai Nguyen et al. [38] | Tomato | Classification | VGG-19 + HSV segmentation | 16,010 images | Accuracy: 99.72% | Dataset from controlled environment |
| Mimma et al. [39] | Fruit | Detection + Classification | YOLOv3/YOLOv7 + ResNet50 | Custom dataset (8 classes) | Accuracy: 99% | Limited fruit categories |
| Elinisa et al. [42] | Banana | Segmentation | U-Net | 18,240 images | Dice: 96.45% | Limited disease classes |

## 3. Methodology

The following discusses the methodology employed to design the automatic system for diagnosing diseases in guava leaves and fruit. Instead of using different processes to classify disease types, detect lesions, and perform pixel-wise segmentation, the methodology uses a consolidated approach that integrates all these processes within a deep learning algorithm. This will be further elaborated on in the next sections discussing the processes employed to prepare and augment the datasets, train the deep learning algorithms, and evaluate their performances. The whole system architecture and its component parts are discussed in the succeeding sub-sections.

### 3.1 Proposed Framework

In particular, the current research makes valuable contributions to the literature on this topic by recognizing several weaknesses in previous studies and developing a deep learning-based framework customized for the automatic identification of diseases associated with guava leaves and fruits. As shown in Fig 1, the proposed framework incorporates three major functionalities, including classification, detection, and segmentation of the pixels. Consequently, all disease symptoms of plants can be analyzed comprehensively within this framework. The whole process involves several stages, including preparation of data set, preprocessing, and augmentation of images, which serve as a solid base for further development of a model. Afterward, the framework continues from disease classification to detection and pixel segmentation. The detection module detects the disease area, and pixel segmentation segments the disease area. The proposed framework improves the understanding of plant disease symptoms. The proposed framework uses explainable AI to highlight areas of concern. The proposed framework uses lightweight and deep models to assess their performance.

**Fig 1.**
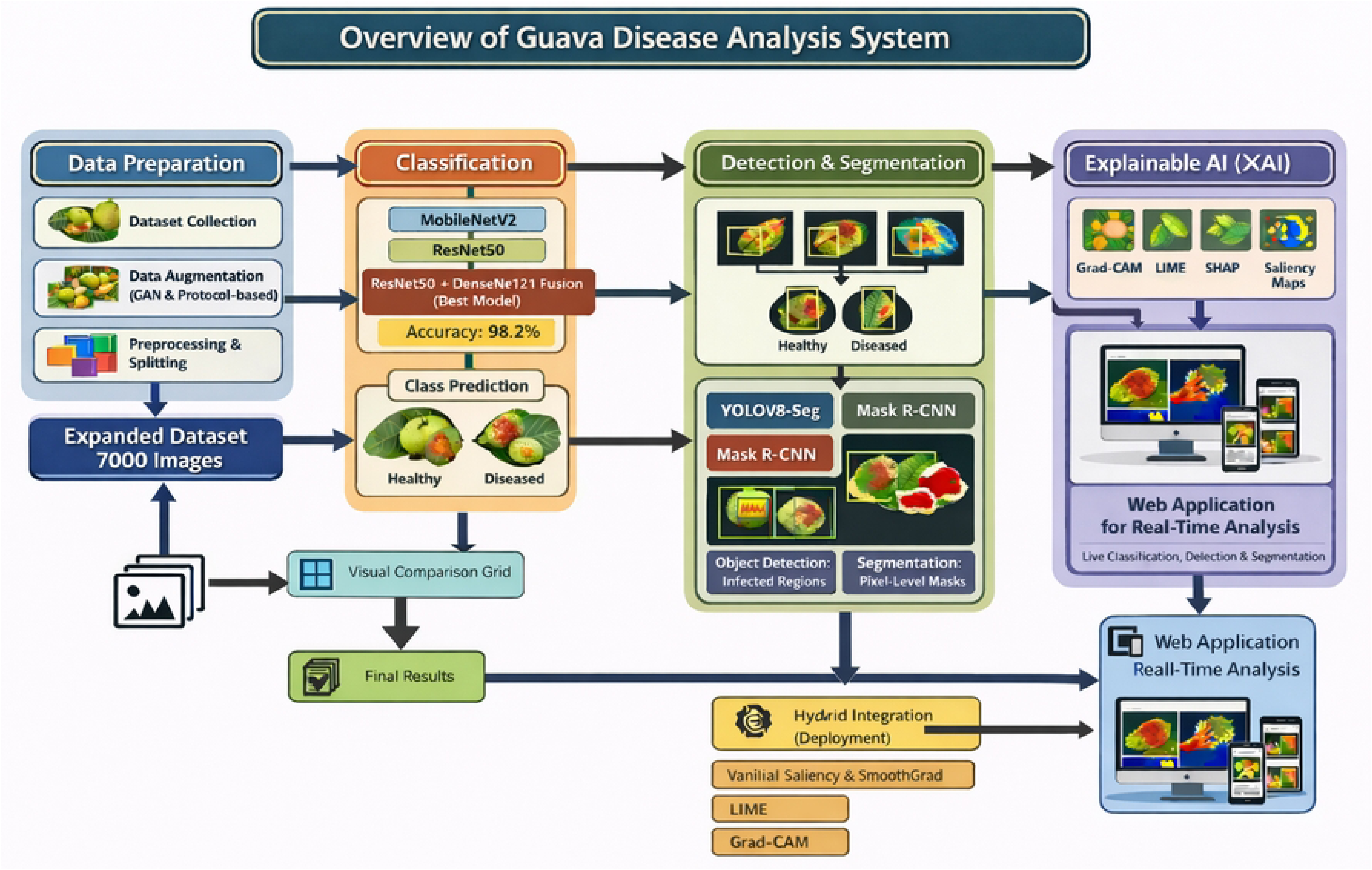
General structure for guava fruit and leaf disease analysis.

### 3.2 Dataset Description and Preprocessing

The dataset used in this research is retrieved from the public access repository on Kaggle, as mentioned in [44], which contains images related to diseases on the leaves and fruits of the guava plant. It comprises 527 images with five different classes: Disease Free (126 images), Phytophthora (114 images), Red Rust (87 images), Styler and Root (94 images), and Scab (106 images). Fig 2 shows the distribution of the images in the dataset, and Fig 3 shows the images for each class.

**Fig 2.**
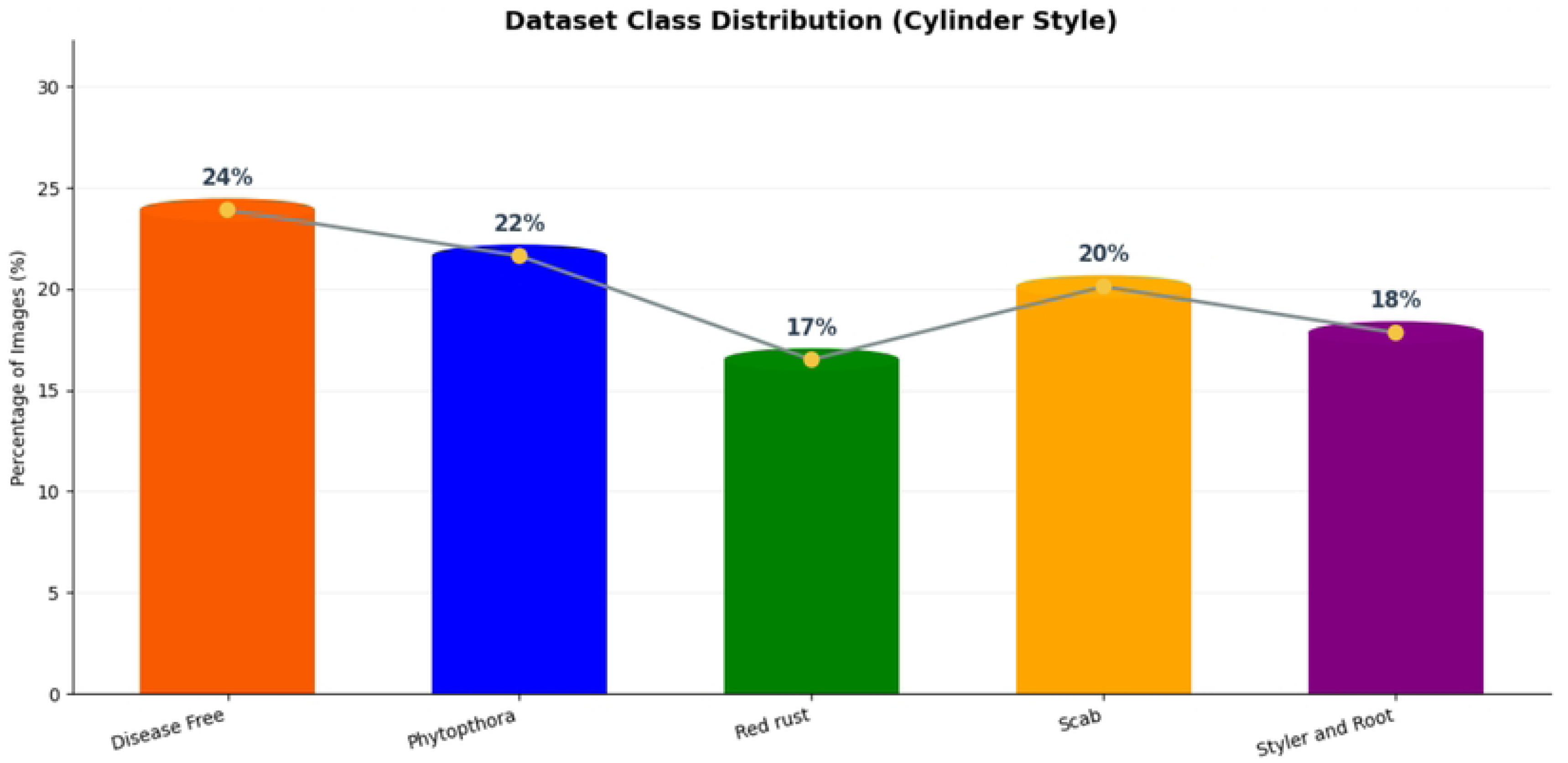
Class distribution of the raw dataset.

**Fig 3.**
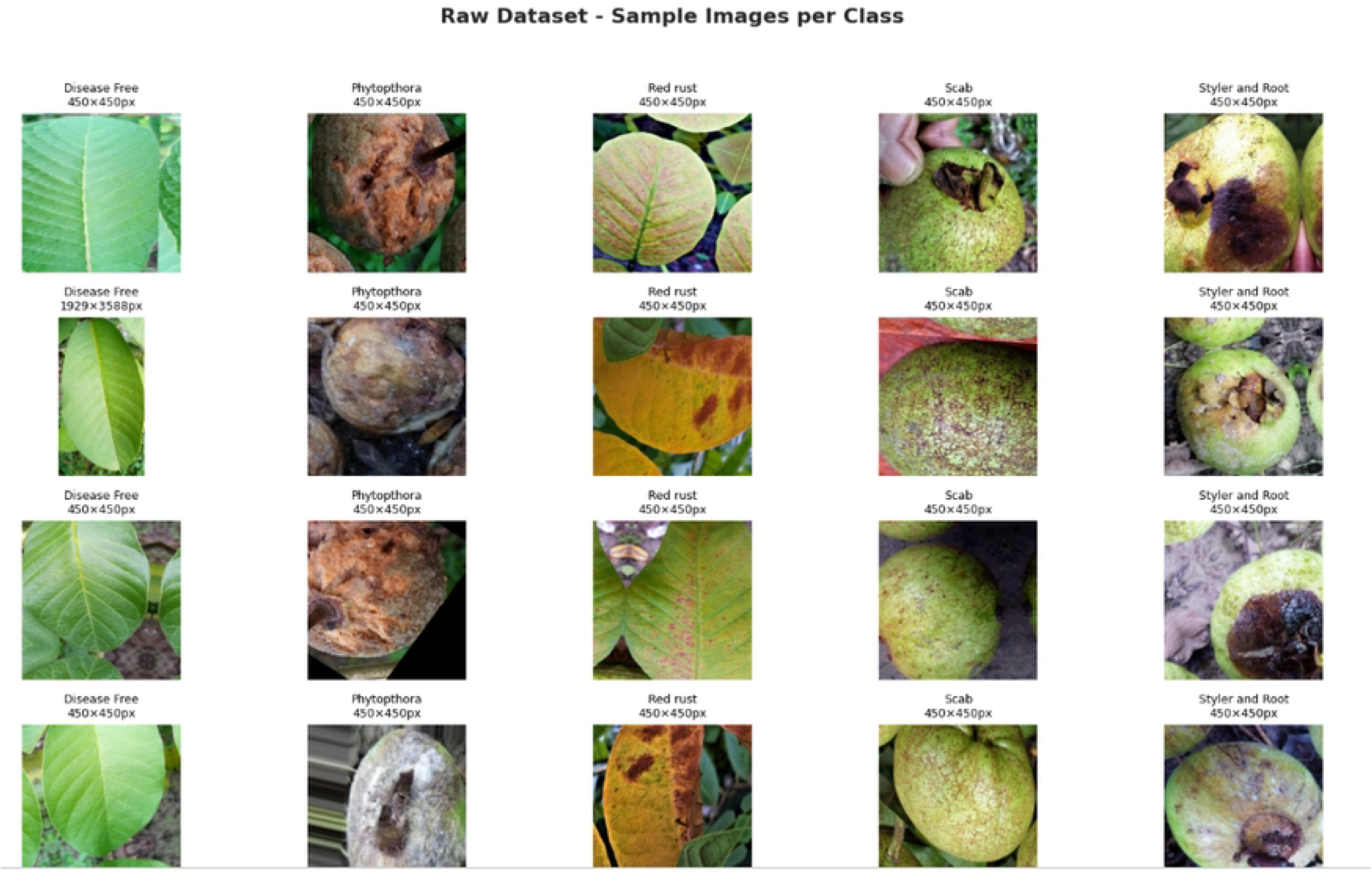
Sample images per class for raw dataset.

The dataset was stratified into training (70%), validation (20%), and testing (10%) sets comprising 369, 105, and 53 images respectively, with the test set strictly held out for final evaluation to prevent data leakage. Within the training set, a stratified 5-fold cross-validation strategy guided model selection and hyperparameter tuning, where augmentation was applied exclusively to the training portion of each fold while fold-validation data remained untouched to prevent data leakage. The validation set additionally monitored training performance and facilitated early stopping for stable convergence. All images were resized to 224×224 pixels, normalized to [0–1], and one-hot encoded. To counter class imbalance, augmentation and oversampling were applied solely to training data, yielding a balanced set of 1,400 images per class, while validation and test distributions remained unchanged to ensure unbiased evaluation. Preprocessing and augmentation examples are illustrated in Figs 4 and 5, with further details summarized in Table 2.

**Fig 4.**
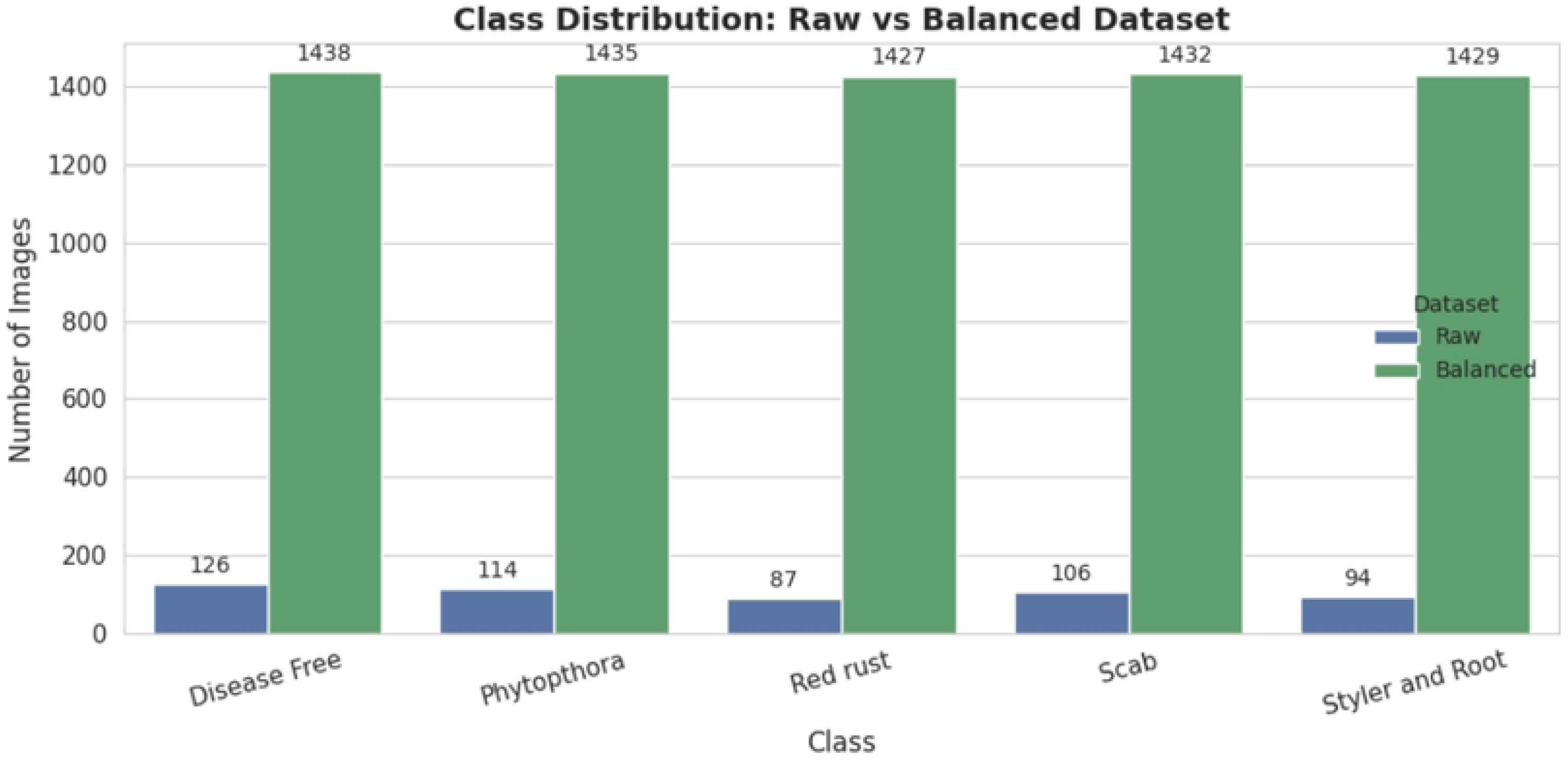
Preprocessing and augmentation comparison of dataset.

**Fig 5.**
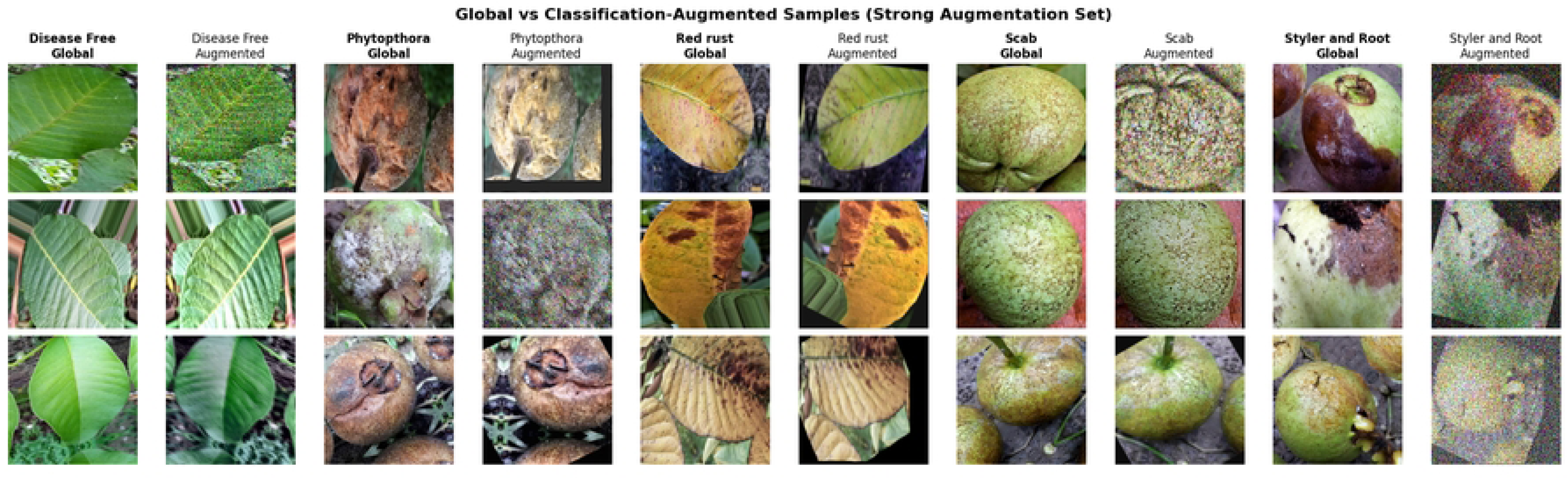
Original vs augmented samples.

**Table 2.** Overview of the dataset at every stage.

| Class | Original | Train (70%) | Validation (20%) | Test (10%) | Augmented Train |
| --- | --- | --- | --- | --- | --- |
| Disease Free | 126 | 88 | 25 | 13 | 1400 |
| Phytophthora | 114 | 80 | 23 | 11 | 1400 |
| Red Rust | 87 | 61 | 17 | 9 | 1400 |
| Styler and Root | 94 | 66 | 19 | 9 | 1400 |
| Scab | 106 | 74 | 21 | 11 | 1400 |
| Total | 527 | 369 | 105 | 53 | 7000 |

### 3.3 Annotation for Detection and Segmentation

All annotations were manually verified to ensure consistency of lesion boundaries across disease classes. The same annotated dataset was used for both YOLOv8-seg and Mask R-CNN to ensure fair comparison under identical labeling conditions. Bounding boxes were used to mark the diseased and healthy regions, while polygon annotations were used to mark the boundaries of the lesions. These processes were done at the pixel level. The images were exported in YOLO format with normalized coordinates. Data augmentation techniques were used. These techniques were implemented using the Albumentations library. Flipping, rotation, scaling, and color changes were applied. These processes were done using the Ultralytics YOLO framework.

### 3.4 Architecture of Applied Models for Classification

This paper used a set of convolutional neural network (CNN) frameworks to detect guava leaf and disease in fruits: DenseNet121, NASNetMobile, MobileNetV2, InceptionV3, Xception, EfficientDualBlock CNN, and a hybrid ResNet50+DenseNet121 model. The choice of these architectures was based on the proven effectiveness in the situation of plant disease recognition and the ability to identify salient visual patterns in agronomic imaging. Transfer learning was used, and the pretrained network weights were fine-tuned in order to boost the generalization performance in an environment with a relatively small set of data.

#### 3.4.1 DenseNet121

DenseNet121 is one model that makes use of dense connectivity, that is, every layer obtains the feature maps of all preceding layers. This architecture allows effective sharing of features and improved gradient propagation, hence allowing the network to remember both the low-level and the high-level visual information. The features make DenseNet121 highly suitable for the classification of plant diseases using moderate-sized databases [45]. The dense connectivity mechanism is defined in Eq (1).

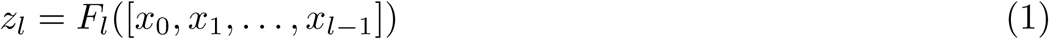

Where, *z_l_* is the output of the *l*th layer and *F_l_*(·) is a composite function that includes convolution, batch normalisation, and nonlinear activation, and [*x*_0_*, x*_1_*, …, x_l−_*_1_] is a concatenation of the feature maps produced by all previous layers.

#### 3.4.2 EfficientDualBlock CNN

Besides pretrained models, a lightweight customized model was developed, which was called EfficientDualBlock CNN, to explore the possibility of a computationally efficient solution to real-time disease classification. It is made up of several dual convolution blocks, which combine standard convolution and depthwise convolution, and then a concatenation of features and pooling layers. The network ends with Global Average Pooling, dense layers, dropout regularization, and a softmax classifier, thus making it possible to efficiently aggregate features with fairly low computational complexity [46]. The convolutional operation that is used in the dual block is mathematically expressed in Eq (2).

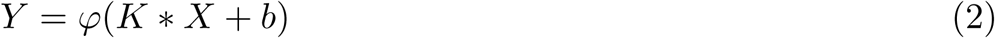

Where, *X* is the input feature map, *K* represents convolution filters, *b* is the bias, ∗ denotes convolution, and *φ* is the activation function.

#### 3.4.3 InceptionV3

Several commonly used CNN architectures were also compared and analyzed. InceptionV3 was chosen because it can extract multi-scale visual features using parallel convolutional filters in its inception modules. This setup allows the network to record disease symptoms with variability in size and texture, a common attribute of plant disease datasets [47]. The output of an Inception module is the resultant activation of the parallel convolutional branches of that module. As shown in Eq (3), features are extracted at different receptive fields across each branch and subsequently concatenated to produce a unified multi-scale feature map.

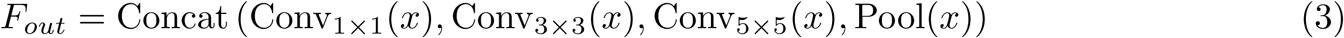

#### 3.4.4 NASNetMobile

NASNetMobile is a lightweight network that is synthesized through Neural Architecture Search (NAS) as it optimally discovers network structures through the use of reinforcement learning methods. The architecture features regular normal cells and reduction cells, hence making it easy to extract features efficiently and maintain low computation complexity that can fit in the mobile platform [48]. The NASNet cell output is depicted in Eq (4).

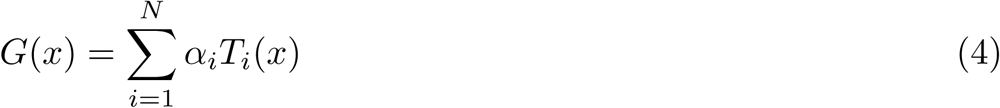

Where, *T_i_*(*x*) denotes candidate operations (e.g., convolution or pooling), and *α_i_* represents their learned weights.

#### 3.4.5 MobileNetV2

MobileNetV2 architecture is designed with computational efficiency via the use of depthwise separable convolutions, inverted residual block, in such a way that there are significant reductions in the number of parameters of the model and floating-point operations, as well as without obvious damage to represent the feature well [48]. Depthwise separable convolution is defined in Eq (5).

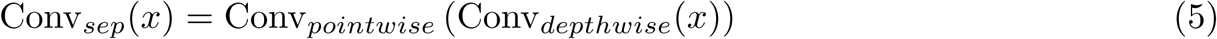

Where, the depthwise convolution is spatial filtering of each channel, and the pointwise convolution (1×1 convolution) is a channel-to-channel information combination.

#### 3.4.6 Xception

The Xception architecture with a more developed variant of this method was also tested due to its better performance in image classification problems that involve complex visual patterns of plant ailments [14]. The convolution operation that is separable and used in the Xception architecture is formally defined in Eq (6).

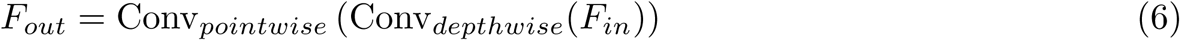

Where, the depthwise convolution is used to obtain spatial features and the pointwise convolution is used to merge information between the channels.

#### 3.4.7 ResNet50+DenseNet121

The fusion model based on ResNet50+DenseNet121 has shown the best classification performance of the tested models, and this is because of the complementary properties of ResNet50+DenseNet121. ResNet50 focuses on abstract structural information using a residual learning approach, whereas DenseNet121 focuses on local details using dense connectivity. In the framework, ResNet50+DenseNet121 are both provided with an input of size 224 × 224 × 3, and their feature maps are downsampled using a Global Average Pooling approach. The feature maps are then integrated to form a single feature vector. The residual learning approach of ResNet50 is given by Eq (7).

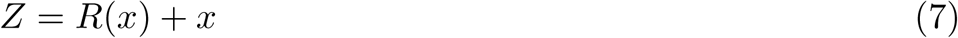

Where, *R*(*x*) represents the residual mapping that is learned by the network. This enables in-depth feature learning while controlling gradient decay. The fused features are then passed through a fully connected layer with 512 neurons, followed by dropout (0.5) and a softmax activation function to obtain probabilities for five different diseases. The overall architecture of the proposed classification model is presented in Fig 6. The complementary features help in improving the classification model. However, the dual backbone nature of this model makes it computationally complex, which makes this model more suitable for server-based agricultural systems than edge-based systems [49]. The training process was carried out using categorical cross-entropy loss, as described in Eq (8), with the Adam optimizer and early stopping, along with a learning rate scheduler, to avoid overfitting.

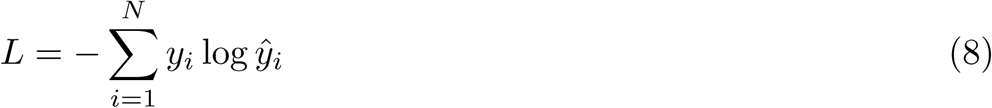

**Fig 6.**
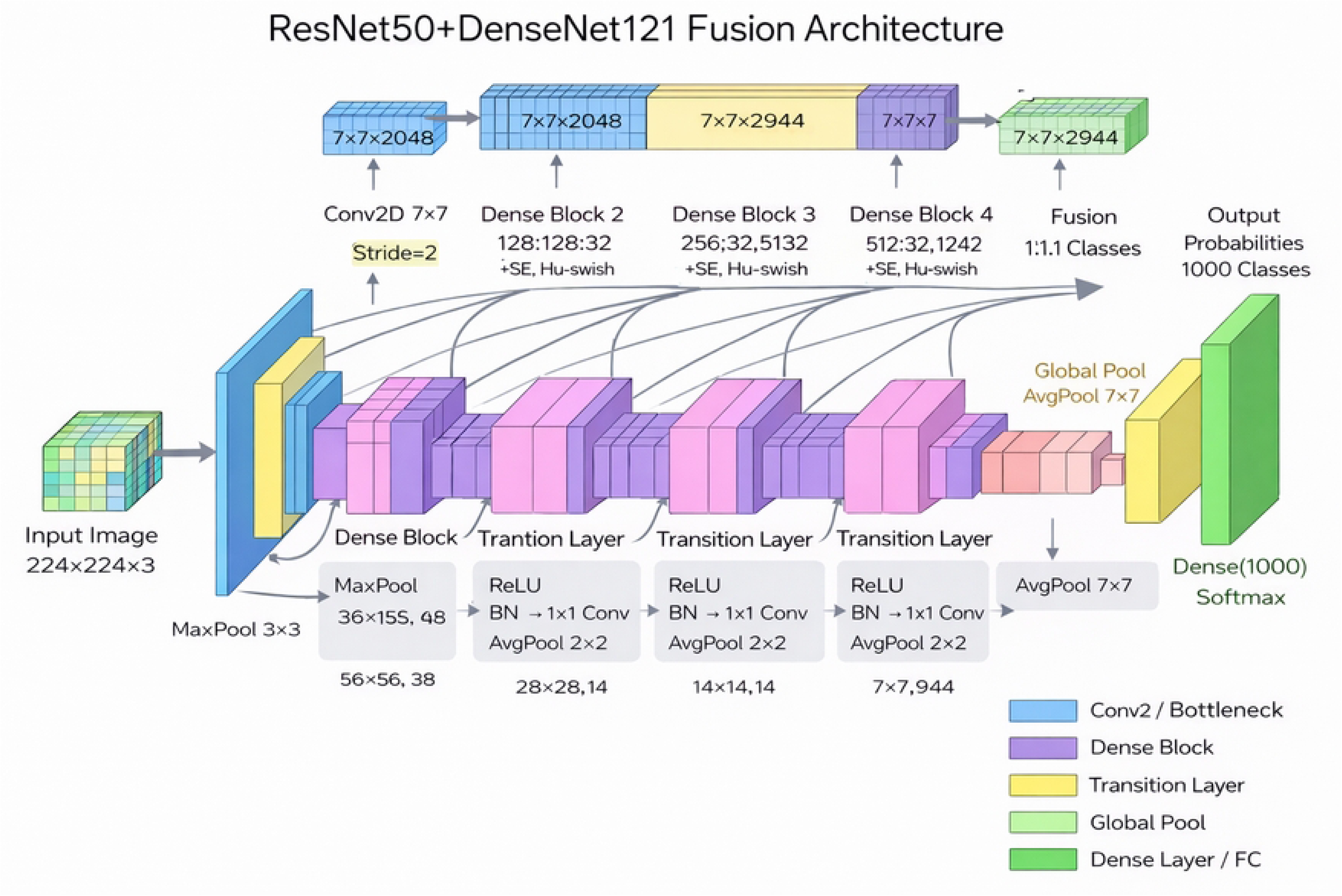
Best model (ResNet50+DenseNet121 fusion) architecture for classification.

### 3.5 Architecture of Utilized Techniques for Detection and Segmentation

#### 3.5.1 Mask R-CNN

The two-stage Mask R-CNN framework has been employed for the identification of diseased regions in guava leaves and fruits. The Mask R-CNN framework utilizes a ResNet50 backbone network in combination with a Feature Pyramid Network (FPN) for hierarchical feature extraction. The FPN enables the ResNet50 network to perform feature extraction on different scales. The Region Proposal Network (RPN) generates candidate regions, which are then aligned using ROIAlign for precise spatial information before classification. Once the diseased regions have been detected, the Mask R-CNN framework performs segmentation on each region by creating masks for each region in parallel. The ResNet50 network has been pre-trained on the COCO dataset, and then the network has been fine-tuned for five different diseases using stochastic gradient descent with data augmentation (7,000 images). The multi-task loss function optimized by the network is given by Eq (9).

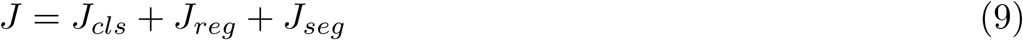

Where, *J_cls_* represents classification loss, *J_reg_* denotes bounding box regression loss, and *J_seg_* corresponds to the segmentation mask loss. The performance of segmentation is measured by calculating the Intersection over Union (IoU) and Dice coefficient, as shown in Eqs (10) and (11). Even though single-stage detectors can perform computations more efficiently, Mask R-CNN allows for more precise localization. Therefore, in order to perform a holistic evaluation of the performance of the detectors, Mask R-CNN and YOLOv8-seg are employed.

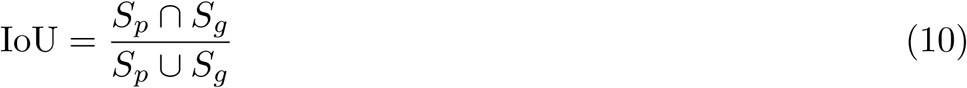

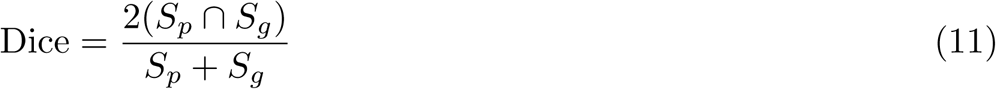

Where, *S_p_*denotes the predicted mask and *S_g_* represents the ground-truth mask.

#### 3.5.2 YOLOv8-seg

The YOLOv8-seg model, part of the Ultralytics library, was utilized to detect and segment infected areas within images of guava leaf and fruit. This model has the advantage of performing object detection and segmentation simultaneously within a single forward pass. This contrasts with other object detection models, where bounding box prediction is performed. The model has an added advantage of predicting bounding boxes and segmentation masks, thus allowing more accurate segmentation of infected areas. The model architecture has a backbone for extracting object features, a neck for performing feature fusion at different scales, and a detection/segmentation head. The backbone layer is responsible for extracting key object features, such as color, texture, and boundary distortions, within infected areas.

The segmentation branch generates a prototype segmentation mask along with coefficients for each infected area. Optimization of the model is performed using a multi-task loss function, as shown in Eq (12).

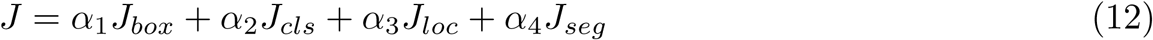

Where, *J_box_*, *J_cls_*, *J_loc_* and *J_seg_* denote bounding box, classification, localization, and segmentation losses, respectively, while *α* are weighting coefficients. The weights were fine-tuned on the labeled dataset. The batch size for training was set to 16, and a total of 100 epochs were carried out. The input resolution for training the network was set to 640 × 640 pixels. Data augmentation methods such as mosaic augmentation, random flipping, and scaling were also carried out for robustness. The overall YOLOv8-seg network is shown in Fig 7.

**Fig 7.**
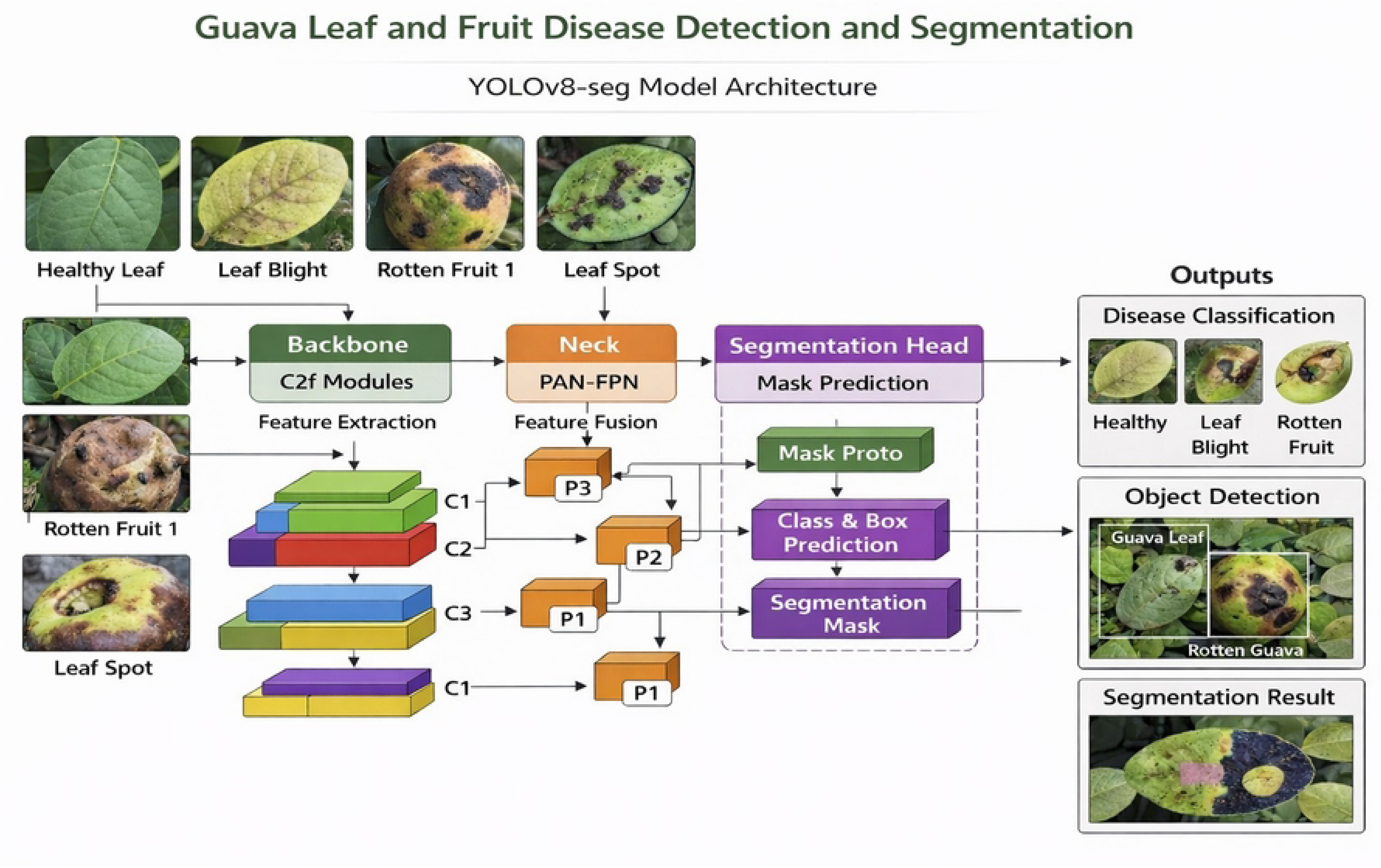
Best model architecture (YOLOv8-Seg) for detection and segmentation.

## 4. Result and Performance Analysis

This section presents the experimental results of the proposed guava disease assessment framework, where all reported metrics reflect final evaluation on the held-out test set, with 5-fold cross-validation confirming consistent performance during model selection. Among the architectures examined, the ResNet50+DenseNet121 fusion model delivered the most reliable classification performance across disease categories, outperforming individual models including DenseNet121 and MobileNetV2, which despite showing good generalization, exhibited residual confusion between visually similar diseases particularly Phytophthora and Styler and Root. ROC, precision-recall, and PCA analyses further validated the fusion model, with most disease classes forming well-separated clusters with minimal overlap. For lesion localization and segmentation, YOLOv8-seg and Mask R-CNN were evaluated for their complementary strengths, where Mask R-CNN demonstrated precise localization and efficient background handling, though its performance declined noticeably for more complex disease categories.

### 4.1 Performance of Applied Models for Classification

Fig 8 shows the confusion matrices for all classification models that have been used for testing in this study. It can be clearly observed from Fig 8(a) that the classification model based on EfficientDualBlock CNN is characterized by the highest level of misclassifications, specifically between categories Styler and Root, and Phytophthora, providing an accuracy rate equal to 90.99%. Gradual improvement can be observed in Fig 8(b) and 8(c) as NASNetMobile and Xception show decreased levels of misclassification with the persistence of certain levels of confusions. In Fig 8(d) and 8(e), further progress can be seen with improved classification rates for InceptionV3 and MobileNetV2 models. Fig 8(f) shows high generalization properties of DenseNet121 with very low levels of confusion, whereas Fig 8(g) illustrates the highest accuracy of 98.20% provided by ResNet50+DenseNet121 fusion.

**Fig 8.**
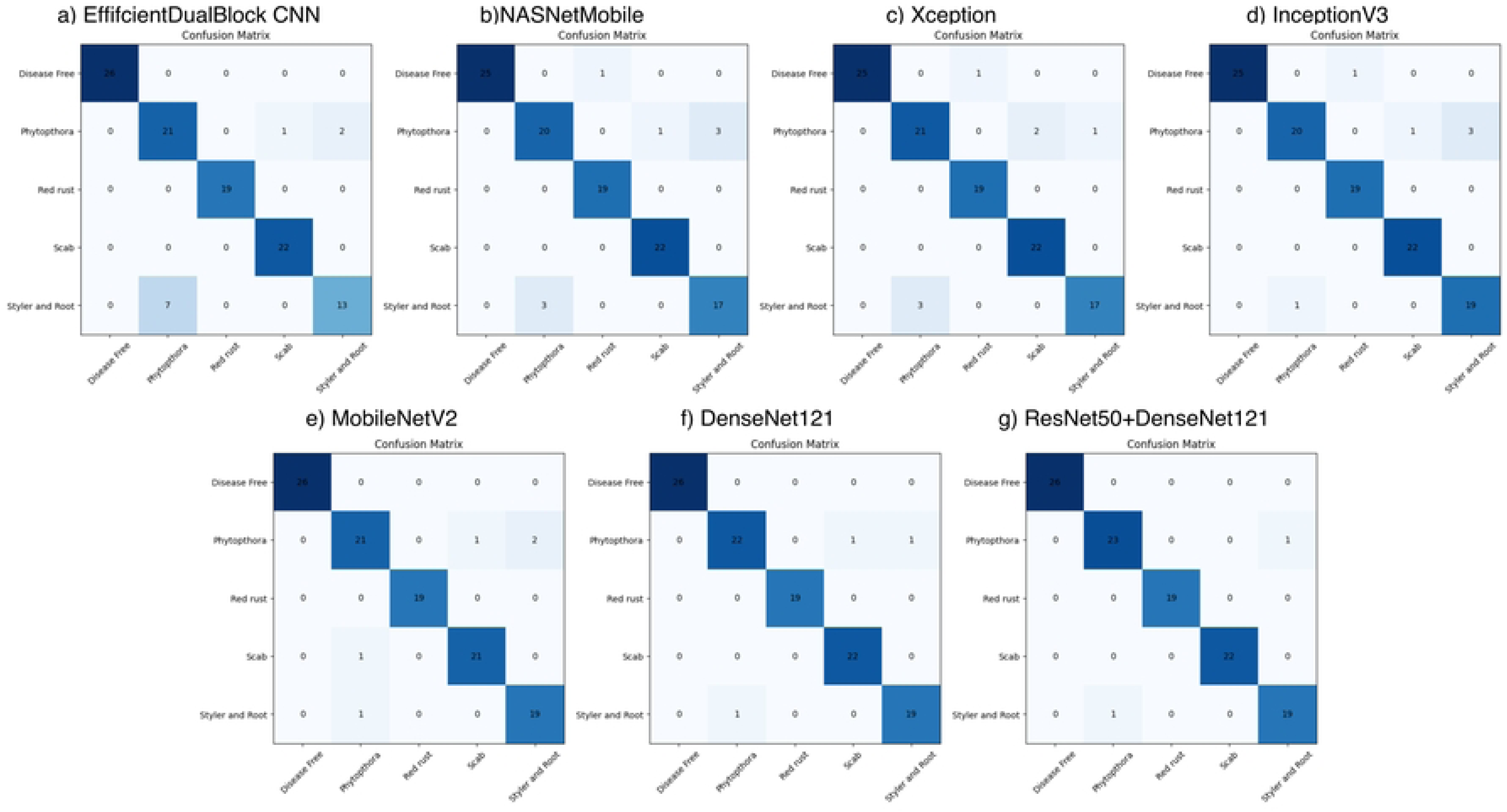
Confusion matrices for the deep learning-based classification models.

To further evaluate classification performance beyond the confusion matrix, ROC curves and AUC values were analyzed to assess the trade-off between TPR and FPR. Fig 9 presents the ROC curves of all models. In Fig 9(a), the EfficientDualBlock CNN shows the lowest performance (AUC 0.975–0.980), with reduced discrimination for Phytophthora (0.950) and Styler and Root (0.972). Performance improves in Fig 9(b) with NASNetMobile (0.991–0.993), though lower AUC remains for Phytophthora (0.973) and Styler and Root (0.986). Fig 9(c) shows Xception (0.997–0.998), followed by Fig 9(d) (InceptionV3, 0.996–0.998) and Fig 9(e) (MobileNetV2, ≈0.996), with slight reductions for similar classes (≈0.991–0.994). Fig 9(f) presents DenseNet121 (≈0.999), achieving 1.0 for Disease Free, Red Rust, and Scab, and 0.998 for Phytophthora and Styler and Root. Finally, Fig 9(g) shows the ResNet50+DenseNet121 fusion model (≈0.999), with 1.0 for Disease Free, Red Rust, and Scab, and near-perfect values for Phytophthora (0.999) and Styler and Root (0.998). Overall, performance improves progressively, while Phytophthora and Styler and Root remain the most challenging classes.

**Fig 9.**
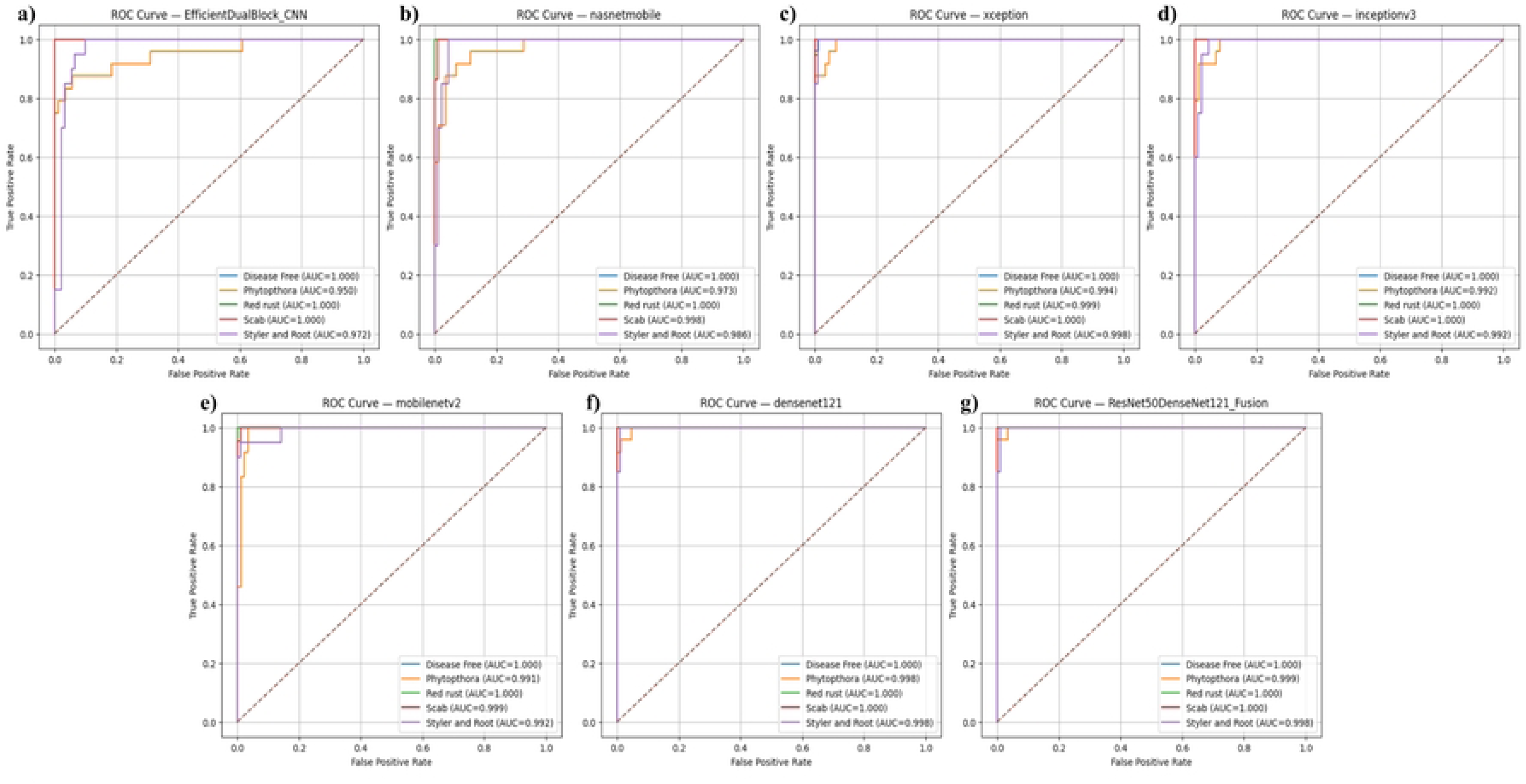
ROC curves of the evaluated deep learning classification models.

To further assess model convergence and generalization, training, validation, and test accuracy-loss curves were examined across all models as illustrated in Fig 10. The EfficientDualBlock CNN in Fig 10(a) exhibits unstable learning characterized by large validation fluctuations (≈80–94%) and elevated loss (≈0.88). Fig 10(b) reveals that NASNetMobile attains high training accuracy (≈99%) but suffers from fluctuating validation performance and higher test loss (≈0.20), suggesting weaker generalization capability. Xception and InceptionV3 in Fig 10(c) and 10(d) demonstrate comparatively stable learning with moderate validation oscillations (≈0.07–0.13) and test accuracy ranging between 93–96%. MobileNetV2 in Fig 10(e) achieves efficient convergence with near-perfect training accuracy and stable validation loss (≈0.05–0.07). DenseNet121 in Fig 10(f) shows strong convergence, maintaining high validation (≈98%) and test accuracy (≈97%) alongside consistently low loss values. The ResNet50+DenseNet121 fusion model in Fig 10(g) delivers the most stable learning overall, recording ≈99.8% training accuracy, ≈98% validation and test accuracy, and remarkably low loss values throughout training (≈0.01), validation (≈0.06–0.08), and testing (≈0.055). Collectively, these curves confirm that deeper and fused architectures yield more stable convergence and superior generalization compared to lighter standalone models.

**Fig 10.**
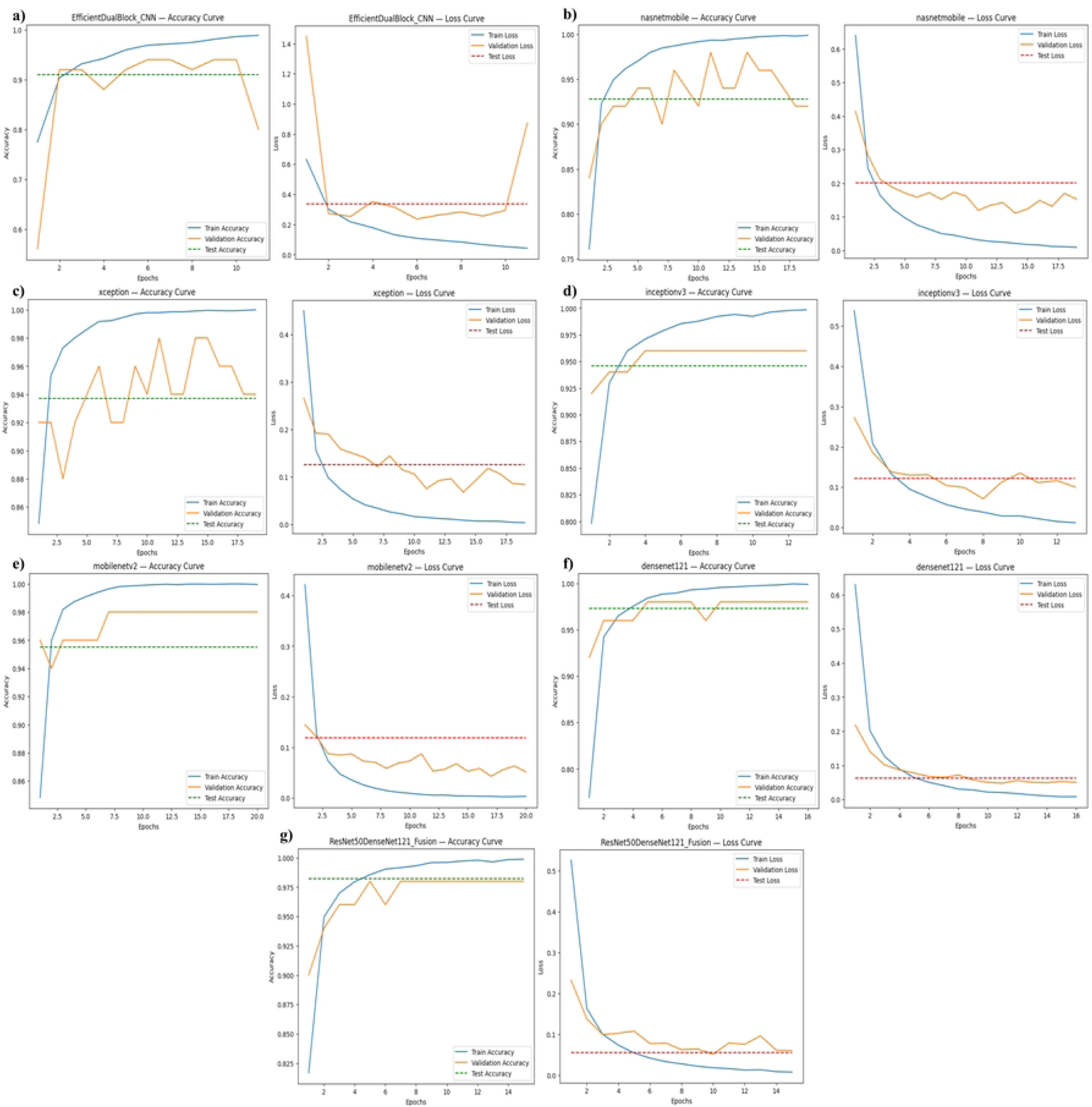
Learning loss and accuracy curves of the evaluated models.

The performance analysis is presented in Table 3, which highlights the performance improvement through the various models. The worst performing model is EfficientDualBlock CNN, since it records the highest loss rate and poor discrimination, especially between classes that are visually alike. However, the performance increases as the models develop from NASNetMobile, Xception, InceptionV3, and MobileNetV2. They are the models that perform the best in terms of accuracy with high AUC value. DenseNet121 records good performance with steady generalization ability and low loss rate. The best performing model is ResNet50+DenseNet121 fusion model. This is evident in Fig 11, which shows the comparison of the average classification metrics among all the models.

**Fig 11.**
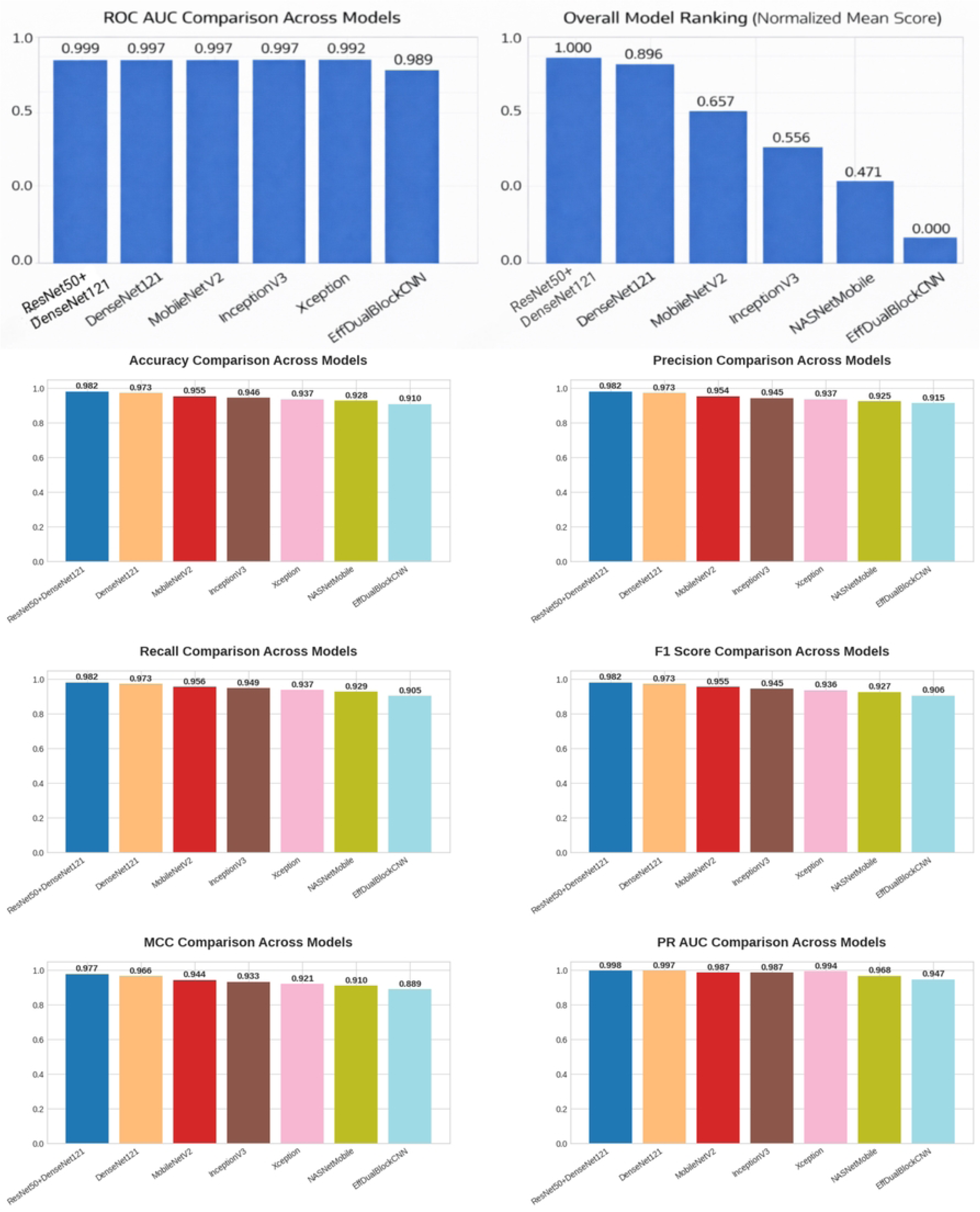
Comparison of the classification metrics.

**Table 3.**
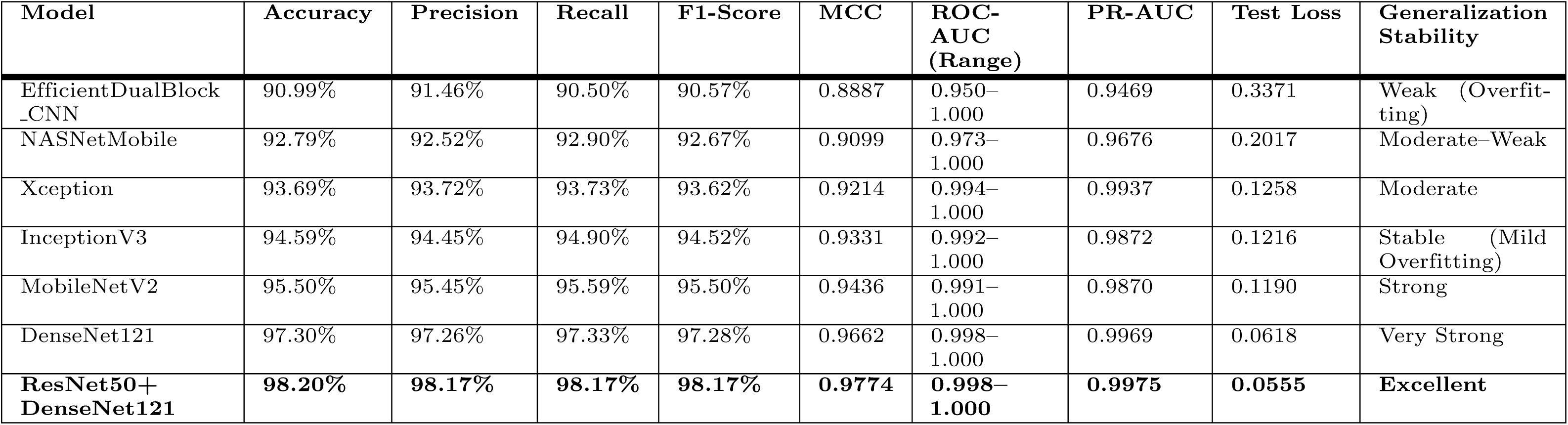
Performance comparison across all applied models.

| Model | Accuracy | Precision | Recall | F1-Score | MCC | ROC-AUC (Range) | PR-AUC | Test Loss | Generalization Stability |
| --- | --- | --- | --- | --- | --- | --- | --- | --- | --- |
| EfficientDualBlock_CNN | 90.99% | 91.46% | 90.50% | 90.57% | 0.8887 | 0.950–1.000 | 0.9469 | 0.3371 | Weak (Overfitting) |
| NASNetMobile | 92.79% | 92.52% | 92.90% | 92.67% | 0.9099 | 0.973–1.000 | 0.9676 | 0.2017 | Moderate–Weak |
| Xception | 93.69% | 93.72% | 93.73% | 93.62% | 0.9214 | 0.994–1.000 | 0.9937 | 0.1258 | Moderate |
| InceptionV3 | 94.59% | 94.45% | 94.90% | 94.52% | 0.9331 | 0.992–1.000 | 0.9872 | 0.1216 | Stable (Mild Overfitting) |
| MobileNetV2 | 95.50% | 95.45% | 95.59% | 95.50% | 0.9436 | 0.991–1.000 | 0.9870 | 0.1190 | Strong |
| DenseNet121 | 97.30% | 97.26% | 97.33% | 97.28% | 0.9662 | 0.998–1.000 | 0.9969 | 0.0618 | Very Strong |
| ResNet50+DenseNet121 | 98.20% | 98.17% | 98.17% | 98.17% | 0.9774 | 0.998–1.000 | 0.9975 | 0.0555 | Excellent |

To improve transparency, a set of explainable AI (XAI) methods, namely Saliency, SmoothGrad, LIME, and SHAP, were applied to the held-out test images to ensure that the predictions were based on disease-relevant regions. The fusion model based on ResNet50+DenseNet121 resulted in an accuracy of 98.20%, an AUC of 0.999, and an MCC of 0.977. The Saliency maps were found to be noisy. SmoothGrad helped improve localization, LIME provided a clear boundary for the lesion, and SHAP provided the most consistent global explanations, which is consistent with the ANOVA results. The overlap between Phytophthora and Styler and Root is due to the similarity in appearance of these two diseases and is consistent with the results obtained from the confusion matrix and PCA. The results of the XAI methods are shown in Fig 12.

**Fig 12.**
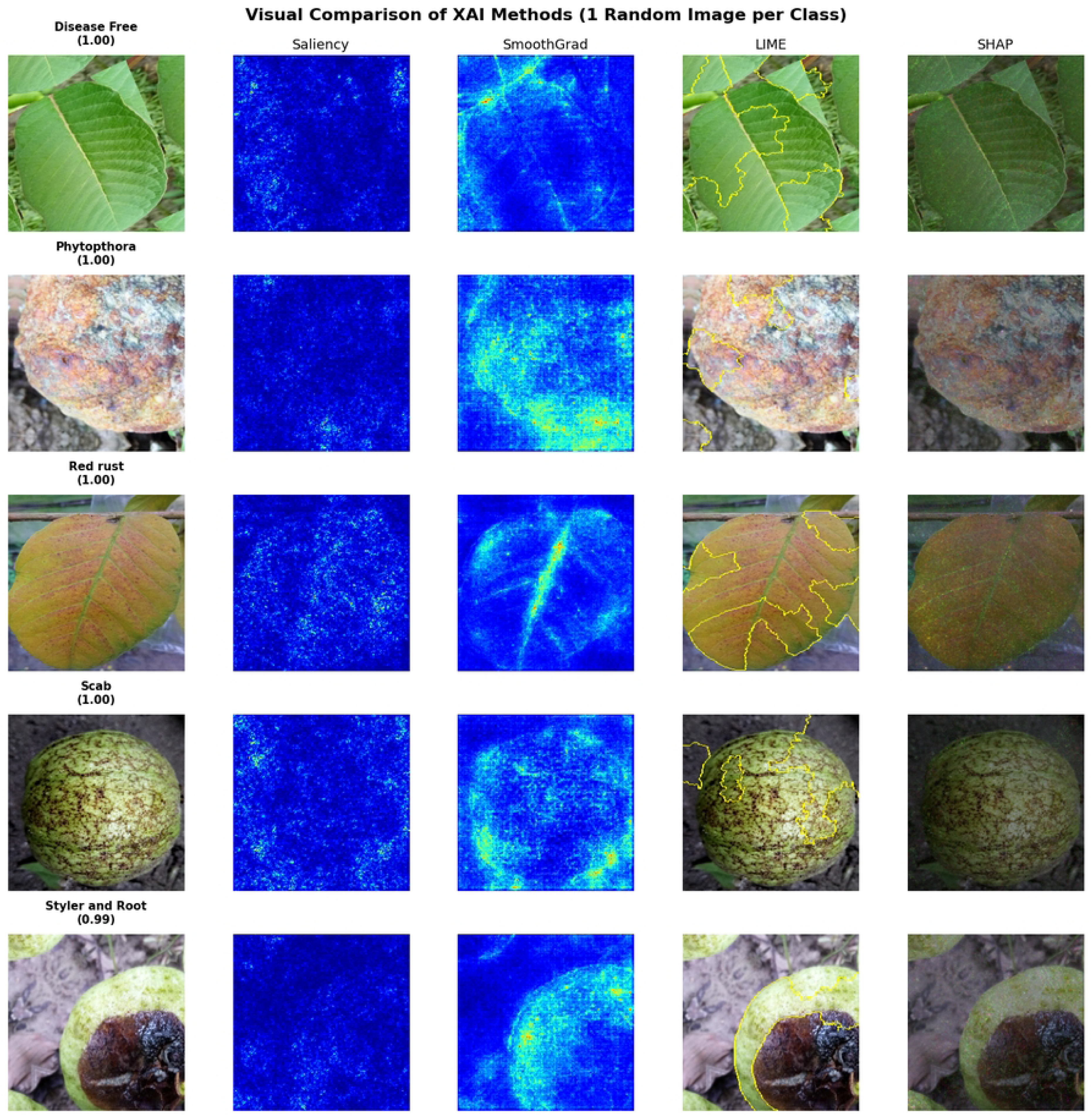
Visual comparison grid (1 random image per class).

### 4.2 Statistical and Feature-Space Analysis

Additional statistical tests were carried out to validate the performance of the classification models. Similarly, feature space evaluation tests were carried out to validate the performance of the feature representations. The tests included McNemar test, ANOVA feature analysis, PCA, and performance comparisons. All these tests helped in achieving a better understanding of the performance of the classification models.

#### 4.2.1 McNemar’s and Chi-square test

The performance of the classification models in terms of prediction was evaluated by means of the McNemar test, which was conducted over the shared test set. Unlike the comparison of accuracy, which only relies on the numerical values, the McNemar test examines the differences based on the patterns of misclassification. Considering the proposed ResNet50+DenseNet121 fusion model as the baseline, there was a statistically significant difference only in the case of EfficientDualBlock CNN, while there were no other significant differences in the performance of the other models. This implies that, in spite of the better performance of the proposed ResNet50+DenseNet121 fusion model in terms of accuracy, the patterns of misclassification of most models are statistically similar in terms of the sample size. The results of the pairwise comparisons can be found in Table 4.

**Table 4.** Pairwise comparisons: ResNet50+DenseNet121 fusion model vs all other models.

| Reference Model | Compared Model | Better Model | McNemar p-value | Significant (p < 0.05) | Chi-square p-value |
| --- | --- | --- | --- | --- | --- |
| ResNet50<br>DenseNet121<br>_Fusion | EfficientDualBlock<br>_CNN | ResNet50<br>DenseNet121<br>_Fusion | 0.0433 | Yes | 0.0209 |
| ResNet50<br>DenseNet121<br>_Fusion | NASNetMobile | ResNet50<br>DenseNet121<br>_Fusion | 0.1138 | No | 0.0578 |
| ResNet50<br>DenseNet121<br>_Fusion | Xception | ResNet50<br>DenseNet121<br>_Fusion | 0.1824 | No | 0.0956 |
| ResNet50<br>DenseNet121<br>_Fusion | MobileNetV2 | ResNet50<br>DenseNet121<br>_Fusion | 0.2482 | No | 0.0833 |
| ResNet50<br>DenseNet121<br>_Fusion | InceptionV3 | ResNet50<br>DenseNet121<br>_Fusion | 0.2888 | No | 0.1573 |
| ResNet50<br>DenseNet121<br>_Fusion | DenseNet121 | ResNet50<br>DenseNet121<br>_Fusion | 1.0000 | No | 0.3173 |

#### 4.2.2 ANOVA Test

To evaluate the discriminative power of learned features, an analysis of variance was performed on learned feature representations using the ResNet50 + DenseNet121 fusion model. The results indicate a strong level of class separability, where multiple features show statistically significant differences across classes of diseases.

This confirms that the model learns informative features to support classification. The distributions of learned features are shown in Fig 13, and a detailed statistic is shown in Table 5.

**Fig 13.**
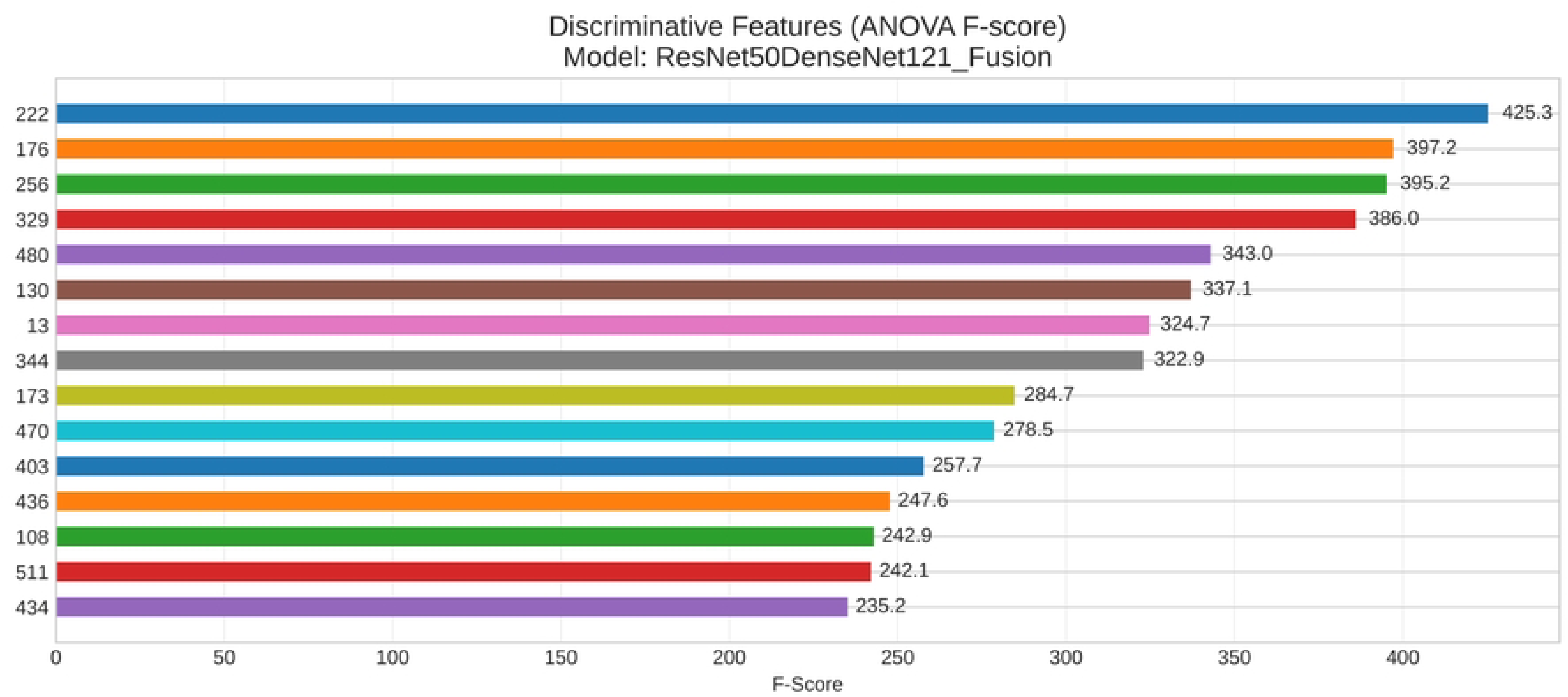
Discriminative features (ANOVA F-score).

**Table 5.**
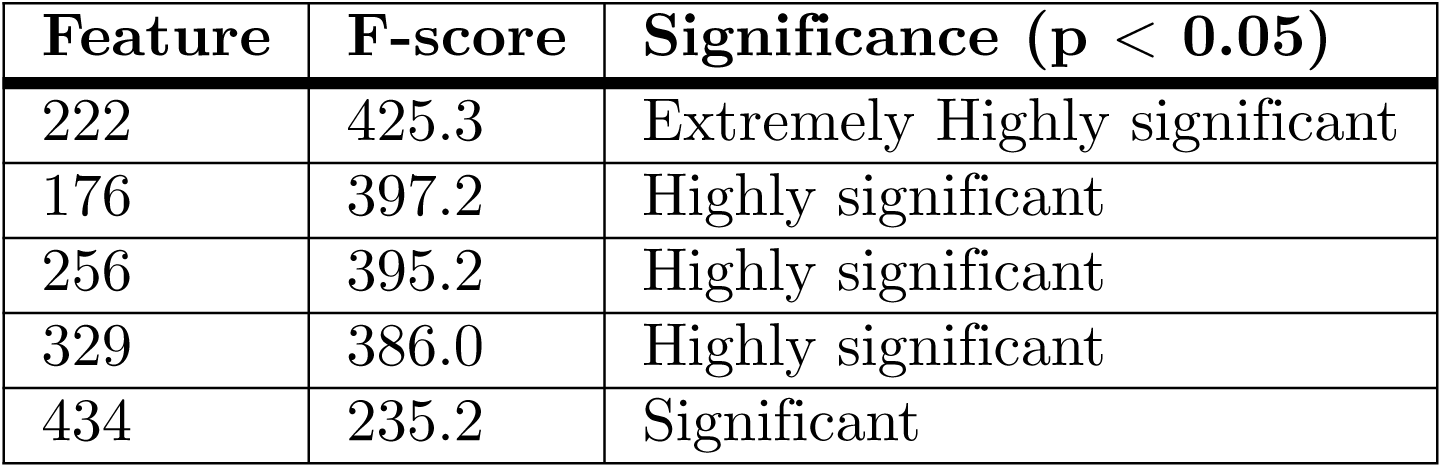
ANOVA test.

#### 4.2.3 PCA Separability

To examine feature separability, Principal Component Analysis (PCA) was applied to the high-dimensional deep features (3,072 dimensions) obtained from the fusion model, combining ResNet50 (≈2,048) and DenseNet121 (≈1,024) features. These embeddings were projected into a two-dimensional space for visualization. The first two components explain 29.3% and 16.4% of the variance, respectively, with a cumulative variance of 45.77%. As shown in Fig 14, most disease classes form distinct clusters: Disease Free and Red Rust are clearly separated, while Scab forms a compact cluster. In contrast, Phytophthora and Styler and Root appear close and partially overlap, consistent with confusion matrix results due to their visual similarity. Overall, the PCA analysis confirms that the fusion model learns discriminative feature representations for effective disease separation.

**Fig 14.**
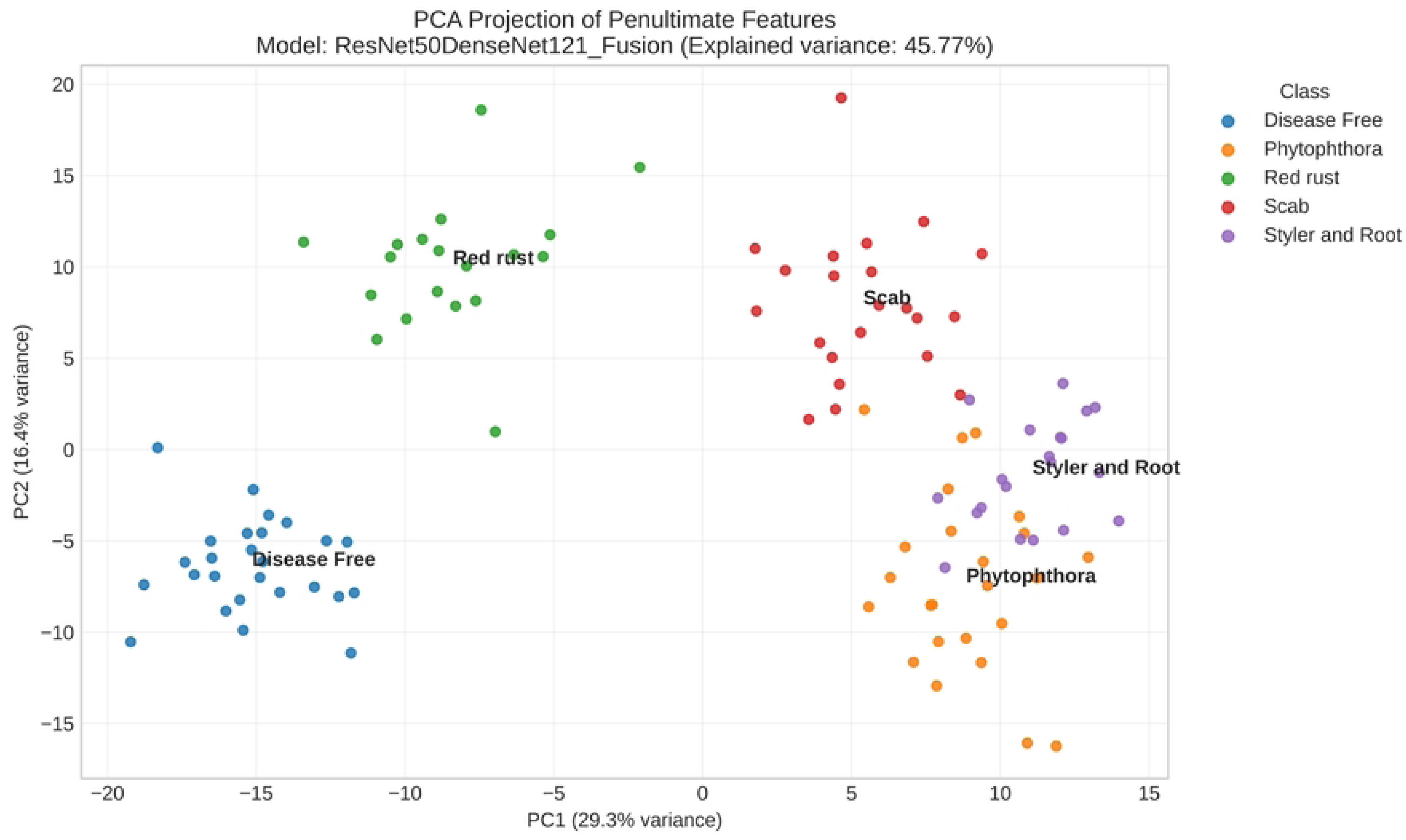
PCA projection of penultimate features.

#### 4.2.4 Performance Comparison (Balanced Accuracy, Specificity, and MCC)

In order to further assess the robustness of the models, Balanced Accuracy, Macro Specificity, and Matthews Correlation Coefficient (MCC) were also assessed. From the results, it is evident that there is consistency in the performance variations among the models. It is also evident that the performance of the ResNet50+DenseNet121 fusion model is the most reliable, while the performance of the EfficientDualBlock CNN is the poorest. A comparative representation of the performance is shown in Fig 15.

**Fig 15.**
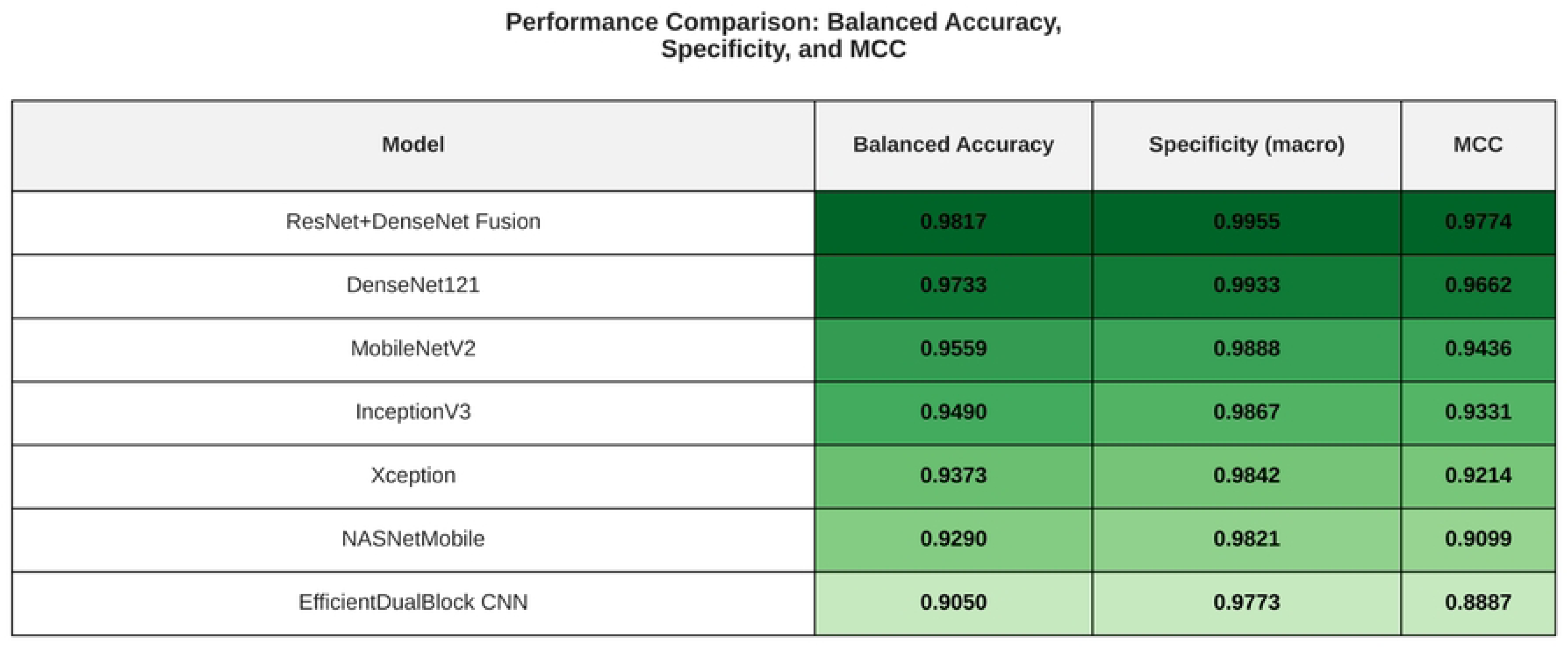
Performance comparison—balanced accuracy, specificity, and MCC.

All in all, the statistical analysis and feature space analysis have shown that the ResNet50+DenseNet121 fusion model is the most reliable and balanced in its classification performance, followed by DenseNet121 and MobileNetV2.

### 4.3 Performance Analysis for Detection and Segmentation

Two advanced architectures, Mask R-CNN and YOLOv8-seg, were evaluated for guava disease localization and segmentation using confusion matrices and detection–segmentation metrics. Mask R-CNN achieved 88.5% accuracy and ≈0.88 balanced accuracy, with strong true-positive predictions, as reflected in Fig 16(a) and Fig 16(c). Disease Free and Red Rust were perfectly classified, while Phytophthora showed moderate performance, and Styler and Root remained the most challenging class. The model maintained a low false-positive rate (11.5%), with a mean IoU of 0.807 and Dice 0.883.

**Fig 16.**
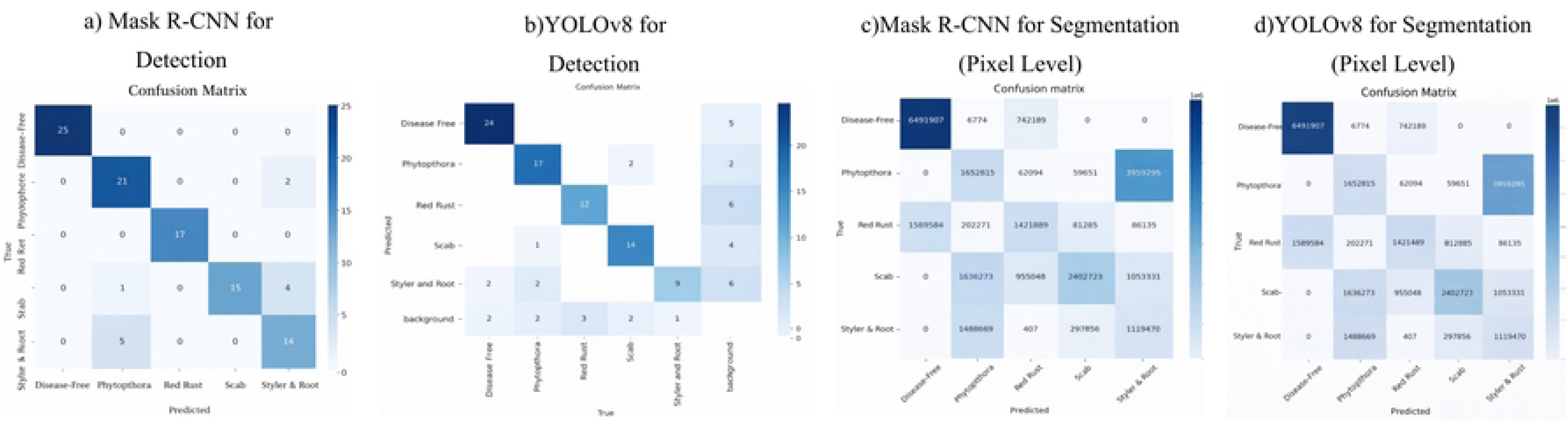
Confusion matrix for detection and segmentation.

In contrast, YOLOv8-seg showed greater inter-class confusion in Fig 16(b) and Fig 16(d), with lower accuracy (66.7%) but similar balanced accuracy (≈0.88–0.89). Despite a higher false-positive rate (24.6%), it achieved superior detection and segmentation performance, with mAP@0.5 of 0.907 and 0.889, and mAP@0.5:0.95 of 0.783 and 0.769, respectively. Detailed quantitative metrics are summarized in Table 6.

**Table 6.** Detection and segmentation metrics.

| Model | Task | mAP@0.5 | mAP@0.5:0.95 | Precision | Recall | F1-score |
| --- | --- | --- | --- | --- | --- | --- |
| **YOLOv8-seg** | Detection (BBox) | **0.907** | **0.783** | 0.899 | **0.881** | **0.890** |
| **Mask R-CNN** | Detection (BBox) | 0.831 | 0.675 | **0.900** | 0.587 | 0.711 |
| **YOLOv8-seg** | Segmentation (Mask) | **0.889** | **0.769** | 0.888 | 0.870 | 0.879 |
| **Mask R-CNN** | Segmentation (Mask) | 0.825 | 0.691 | **0.896** | **0.880** | **0.883** |

Fig 17 shows the class-wise precision–recall (PR) curves evaluated at an IoU threshold of 0.5. For YOLOv8-seg, AP is highest for Disease Free (0.924), followed by Scab (0.911) and Phytophthora (0.887), while lower values for Styler and Root (0.849) and Red Rust (0.800) indicate greater difficulty due to visual similarity. The overall performance reaches mAP@0.5 = 0.874, reflecting a balanced precision–recall trade-off with a typical precision drop at higher recall.

**Fig 17.**
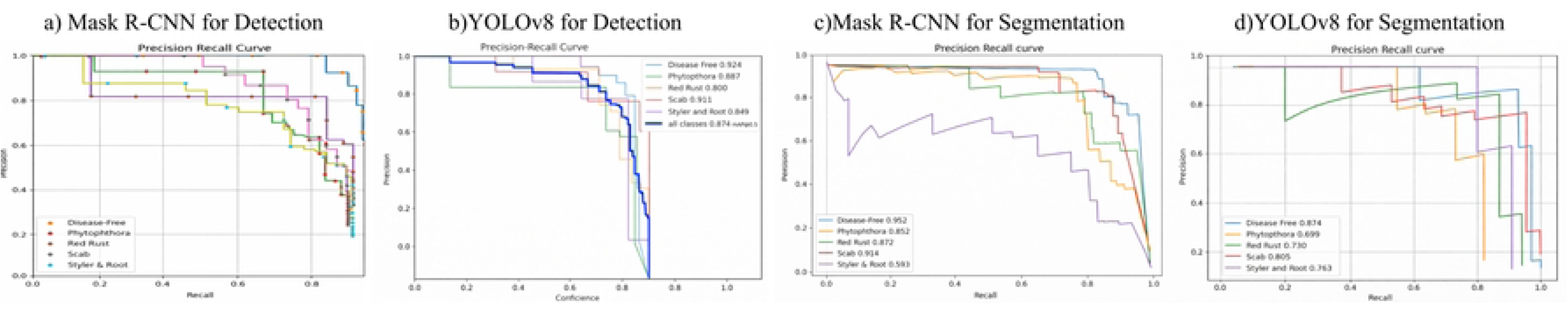
The precision-recall (PR) curves.

For Mask R-CNN, PR curves remain stable for distinct classes (Disease Free, Scab) but show reduced precision for Phytophthora, Red Rust, and Styler and Root at higher recall. Overall, YOLOv8-seg performs better on challenging classes, while Mask R-CNN maintains stronger precision for well-defined categories.

Fig 18 shows the F1–confidence curves for both models. YOLOv8-seg achieves a peak mean F1-score of ≈0.89 at a confidence of ≈0.65, with Disease Free exceeding 0.9. Phytophthora, Red Rust, and Scab show moderate performance, while Styler and Root remain the most challenging. Most classes maintain F1 *>* 0.8 across a wide confidence range (≈0.4–0.8), indicating stable performance.

**Fig 18.**
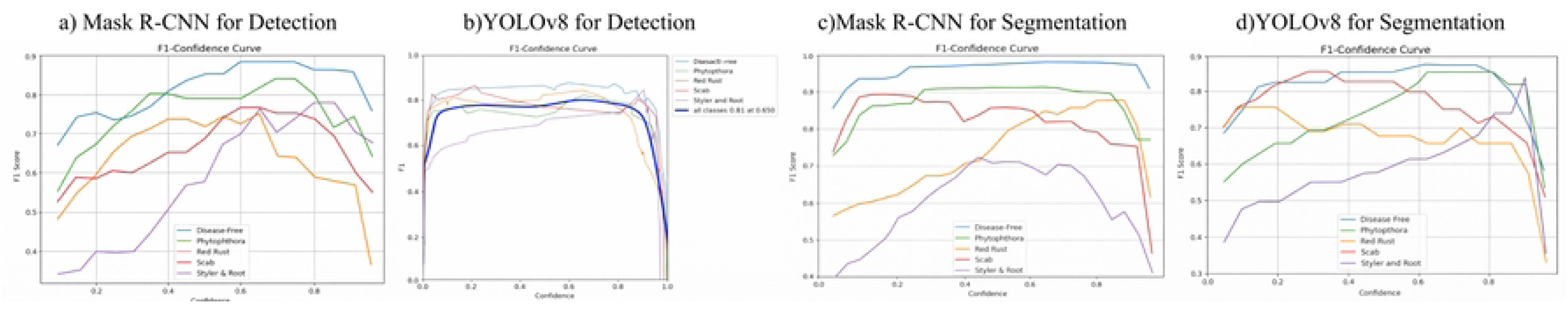
F1-confidence curves.

In contrast, Mask R-CNN reaches a lower peak F1-score (≈0.81) with a narrower stable range. Overall, YOLOv8-seg demonstrates higher and more consistent F1 performance across confidence levels.

Despite strong performance, YOLOv8-seg remains a black-box model, making interpretability essential. To address this, Saliency, SmoothGrad, LIME, and SHAP were applied to highlight pixel-level contributions and ensure predictions focus on lesion regions. YOLOv8-seg outperformed Mask R-CNN in detection and segmentation, achieving mAP@0.5/0.5:0.95 of 0.907/0.783 and 0.889/0.769, compared to 0.831/0.675 and 0.825/0.691, respectively, along with higher recall (0.881 vs. 0.587) and F1-score (0.890 vs. 0.711). As shown in Fig 19, Saliency produces noisy maps, SmoothGrad reduces noise, LIME yields fragmented regions, and SHAP provides the most coherent and semantically meaningful explanations. Overall, explanation quality ranks SHAP *>* SmoothGrad *>* LIME *>* Saliency.

**Fig 19.**
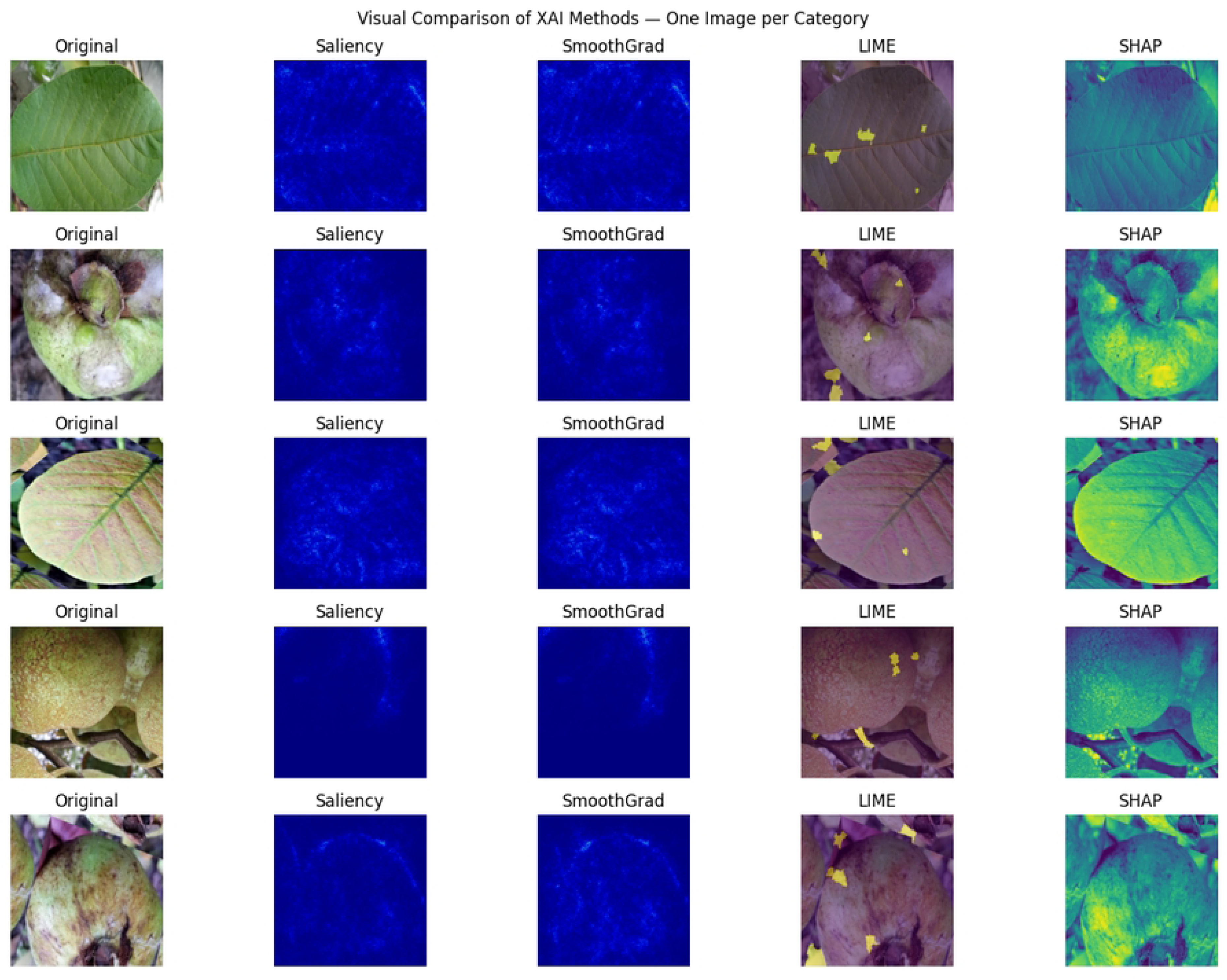
Visual comparison grid (1 random image per class).

Overall, both the classification fusion model and YOLOv8-seg framework are based on biologically significant disease features, which can be considered as a measure of transparency, accuracy, and feasibility of the proposed deep learning framework in diagnosing the disease in guavas.

### 4.4 Deployment and Web-Based Implementation

In the following section, the process of developing the web-based decision support system, GuavaVision AI, is discussed. This system is specifically designed to help in the automated detection of diseases that occur in guava fruits as well as leaves. The system is designed such that the classification, detection, and segmentation techniques that are powered by deep learning can be performed together in one single pipeline, thus helping in the real-time diagnosis of diseases through the client-server architecture.

#### 4.4.1 System Architecture

The GuavaVision AI system is based on a client-server architecture, which is built on a React-based client and a Flask server that provides a RESTful API and cross-origin resource sharing (CORS) for efficient communication. The inputs provided by the client in the form of guava images are Base64 encoded and sent over an HTTP request. The inputs are processed and converted into a final output by decoding the inputs, resizing them, normalizing them, and reconstructing them into RGB form. The unified inference pipeline is based on a fusion model (ResNet50+DenseNet121) for classification and YOLOv8-seg for detection, segmentation, and localization. The output of the inference is provided in a structured JSON format and is also shown to the client in the form of a canvas overlay. The architecture is based on TensorFlow and Keras for a GPU-enabled Flask server. The overall architecture is shown in Fig 20.

**Fig 20.**
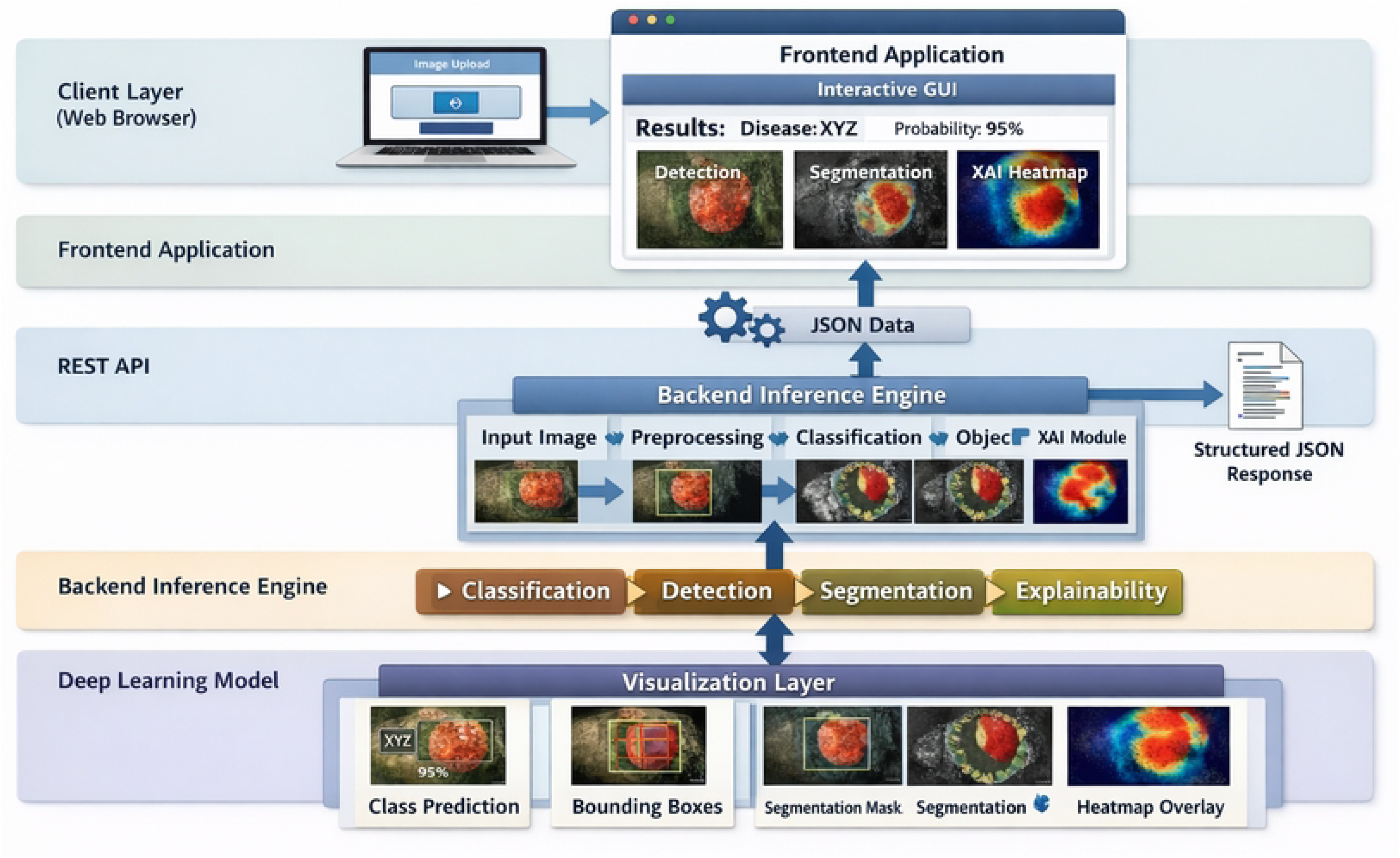
Overall software architecture of the GuavaVision AI web application.

#### 4.4.2 User Interface and System Demonstration

The GuavaVision AI system has been implemented and deployed as a web-based application, which can be accessed through https://guavavision-ai.vercel.app/. The system has a simple and intuitive interface, which helps in real-time interaction with the underlying deep learning models.

The user interface, as represented in Fig 21(a), helps users in uploading images of guava leaves or fruits using a drag-and-drop option or by selecting images from their local machine. Once the images are submitted, the system classifies them and displays the predicted disease along with confidence levels, as represented in Fig 21(b). Further, the system detects the areas of infection using bounding boxes, as represented in Fig 21(c), and segments the images to highlight areas of infection, as represented in Fig 21(d).

**Fig 21.**
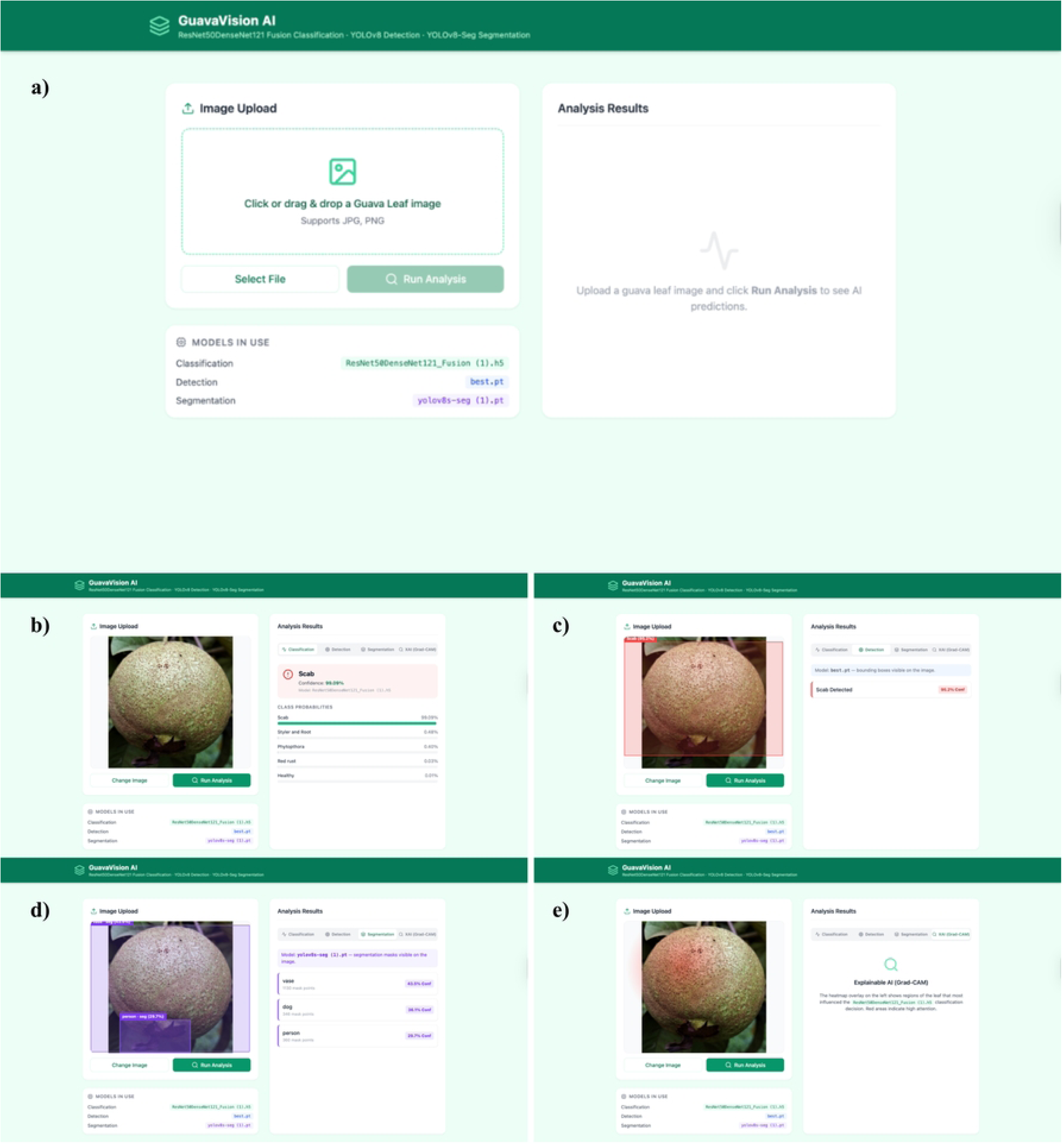
User interface and system demonstration of GuavaVision AI.

To help users interpret the results, the system offers explainable AI results through Grad-CAM, which helps in identifying areas with the strongest influence on the predicted result, as represented in Fig 21(e). This helps users understand both the predicted disease and areas of infection on the guava leaves, which in turn helps in decision-making.

## Conclusion

This study presented a unified deep learning framework for the classification, detection, and pixel-level segmentation of guava leaf and fruit diseases. Among the evaluated classification models, the ResNet50+DenseNet121 fusion model achieved the best performance (98.20% accuracy), which showed that the fusion of residual learning with dense feature reuse has a significant impact on multi-class classification of plant diseases. DenseNet121 performed well independently as well, and the performance of MobileNetV2 was an efficient lightweight option that could be deployed to the web.

To localize and segment diseases, YOLOv8-seg was found to perform better than Mask R-CNN with higher detection and segmentation accuracy (mAP@0.5 = 0.907 and 0.889) and faster inference, which makes it an appropriate choice in real-time agricultural monitoring. The interpretability of the models was confirmed through Grad-CAM, Saliency, SmoothGrad, LIME, and SHAP, and regularly demonstrated the presence of disease lesions and patterns of infections, which confirms biologically significant predictions. The experimental results demonstrate that integrating classification, detection, and segmentation within a unified framework improves both diagnostic accuracy and interpretability, making the system suitable for practical agricultural applications.

Although the results were encouraging, the visual similarity between some of the diseases (e.g., Phytophthora and Styler and Root) as well as the relatively small sample size (527 images) can restrict the generalization. In the future, the direction of work involves increased field datasets, lightweight edge portable models, transformer-based architectures, and translation across crop diseases to create a further advance in AI-centered precision agriculture.

